# Evidence for phospholipid export from the gram-negative inner membrane: time to rethink the Mla pathway?

**DOI:** 10.1101/388546

**Authors:** Gareth W. Hughes, Stephen C. L. Hall, Claire S. Laxton, Pooja Sridhar, Amirul H. Mahadi, Caitlin Hatton, Thomas J. Piggot, Mohammed Jamshad, Vaclav Spana, Cadby T. Ian, Christopher Harding, Georgia L. Isom, Jack Bryant, Rebecca J. Parr, Yasin Yakub, Mark Jeeves, Damon Huber, Henderson R. Ian, Luke A. Clifton, Lovering L. Andrew, Timothy J. Knowles

## Abstract

The Mla pathway is believed to be involved in maintaining the asymmetrical Gram-negative outer membrane via retrograde phospholipid transport. The pathway is composed of 3 components: the outer membrane MlaA-OmpC/F complex, a soluble periplasmic protein, MlaC, and the inner membrane ATPase, MlaFEDB complex. Here we solve the crystal structure of MlaC in its phospholipid free closed apo conformation, revealing a novel pivoting β-sheet mechanism which functions to open and close the phospholipid-binding pocket. Using the apo form of MlaC we provide evidence that the Mla pathway functions in an anterograde rather than a retrograde direction by showing the inner membrane MlaFEDB machinery exports phospholipids and transfers them to MlaC in the periplasm. We confirm that the lipid export process occurs through the MlaD component of the MlaFEDB complex. This lipid export process is shown to be independent of ATP. Our data provides, for the first time, evidence of an apparatus for lipid export to the outer membrane.

## Introduction

Rising antimicrobial resistance, particularly in difficult-to-treat Gram negative infections, is rapidly becoming one of the biggest challenges facing the 21^st^ century. A key research priority now lies in identifying conserved pathways in Gram-negative bacteria that could be exploited as novel drug targets (WHO 2017). Of particular interest are the pathways involved in the biosynthesis of the Gram-negative cell envelope due to their widespread conservation and proximal location to the bacteria surface.

Gram-negative bacteria have a complex envelope consisting of an inner (IM) and outer membrane (OM), separated by a thin peptidoglycan layer and aqueous periplasm (Malinverni and Silhavy 2009). The OM has a unique structure and contains two types of lipids, phospholipids (PL) and lipopolysaccharide (LPS), which are asymmetrically distributed. PL is found primarily in the inner leaflet and LPS is found in the outer leaflet. The membrane also harbours unique proteins, known as outer membrane proteins that contain a β-barrel architecture, as well as containing lipoproteins which are predominantly found at the inner leaflet. The structure of this asymmetric membrane serves as a formidable barrier against environmental stress, antibiotics and immune factors (Ruiz, Kahne et al. 2006). The asymmetry is thought to be a key factor in maintaining barrier function as previous studies have shown that PLs are only found in the outer leaflet under stress conditions leading to reduced OM integrity (Kamio and Nikaido 1976, Nikaido 2005).

Research from various groups over the last two decades has provided considerable mechanistic understanding of the processes involved in the formation and trafficking of components to the OM, including the β-barrel assembly machine (Bam) complex, periplasmic chaperones (Wu, Malinverni et al. 2005, Malinverni, Werner et al. 2006, Sklar, Wu et al. 2007, Sklar, Wu et al. 2007, Knowles, 2009 #145), the Lpt pathway for LPS transport (Ruiz, Gronenberg et al. 2008, Narita and Tokuda 2009, Chng, Gronenberg et al. 2010, Tran, Dong et al. 2010) and the Lol pathway for lipoprotein trafficking to the OM (Okuda and Tokuda 2011, Grabowicz and Silhavy 2017). In the case of PLs, it is known there is bidirectional movement of PLs between the IM and OM, (Jones and Osborn 1977, Langley, Hawrot et al. 1982). Furthermore, through pulse-chase experiments it was demonstrated that PL translocation can occur rapidly and independently of ATP hydrolysis (Donohue-Rolfe and Schaechter 1980). However, the mechanism by which PLs are trafficked between the membranes remains largely unresolved.

The first evidence for a proteinaceous system responsible for PL trafficking between the IM and OM came in the form of the Maintenance of Lipid Asymmetry (Mla) pathway, a six protein (MlaA-F) system with components in the IM, periplasm, and OM. It was described as having a role in maintaining lipid asymmetry in the OM by removing and trafficking PLs found in the outer leaflet of the OM to the IM (Malinverni and Silhavy 2009). Through genetic knockouts, structural and modelling-based studies, the individual components have been identified and characterised (Figure 1)(Chong, Woo et al. 2015, Thong, Ercan et al. 2016, Abellon-Ruiz, Kaptan et al. 2017, Ekiert, Bhabha et al. 2017, Yeow, Tan et al. 2018). Furthermore, several studies have indicated that they are important for infection (Suzuki, Murai et al. 1994, Hong, Gleason et al. 1998, Cuccui, Easton et al. 2007). The components include an OM lipoprotein termed MlaA, which has been found to be associated with the major porins OmpF and OmpC, and is believed to extract PLs from the OM, a soluble periplasmic component known as MlaC, thought to ferry PLs between the membranes; and an inner membrane ATPase composed of MlaF, MlaE, MlaD and MlaB in the following stoichiometry’s 2:2:6:2 that is assumed to accept and insert PLs into the IM. However, little information regarding the mechanism of this proposed PL trafficking pathway has been confirmed. In particular, questions still remain regarding the directionality of PL transport, and why such a complex transport mechanism is present when other simpler systems, such as the OM phospholipase A2, PldA (Snijder, Ubarretxena-Belandia et al. 1999) and the lipopolysaccharide palmitoyl transferase, PagP (Bishop, Gibbons et al. 2000), are available in the outer membrane to maintain asymmetry?

**Figure 1:**
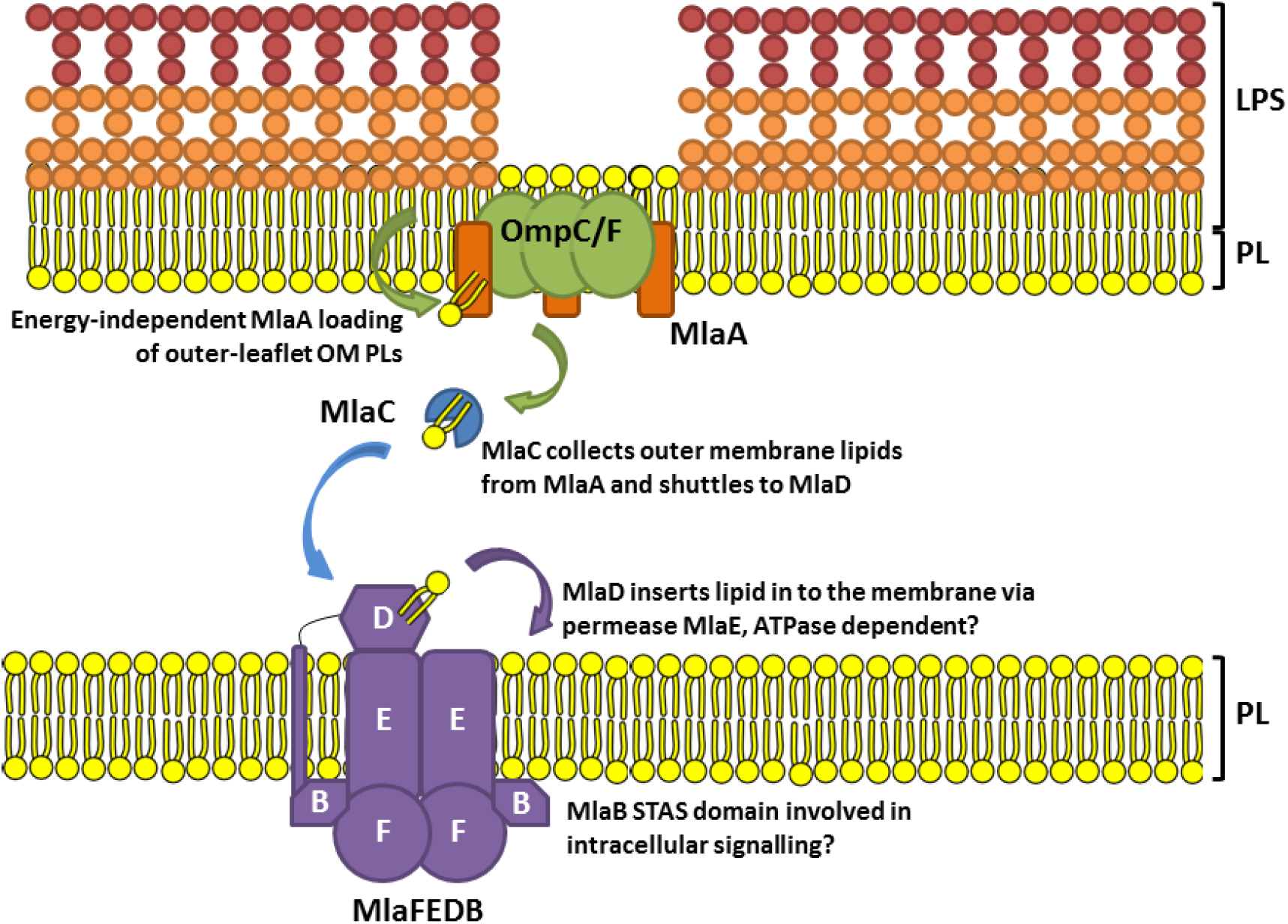
**Schematic of summary of the current proposed Mla pathway mechanism**, based on work by Malinverni & Silhavy, 2009, Ekiert *et al*. 2017 and Abellón-Ruiz *et al*. 2017. The Mla pathway is thought to constitute a retrograde PL transport system for the removal of misplaced OM PLs, in which MlaA removes OM PLs and passes them to MlaC, which shuttles them through the periplasm to MlaFEDB for reinsertion into the IM (May and Silhavy 2017). The MlaFEDB complex is a classic ABC transporter, with MlaE being the integral membrane protein and predicted permease (Thong, Ercan et al. 2016).

In this study we sought to address these questions focusing on the interaction between the periplasmic MlaC and the MlaFEDB complex. Here we provide evidence that the MlaFEDB complex functions in an anterograde direction, extracting PLs from the IM and passing them to MlaC via MlaD, and that it does so in an ATP independent manner. Furthermore, we determine the 2.25Å crystal structure of the apo form of MlaC and reveal that it functions by a novel β-pivot mechanism that requires interaction with MlaFEDB for PL loading. This information together provides the first evidence of a pathway involved in anterograde PL transport towards the OM.

## Methods

### Expression and purification of MlaC and MlaD

DNA corresponding to MlaC was previously cloned in to a custom plasmid (pBE1203) with an N-terminal 6x His tag followed by a TEV protease cleavage site (Ekiert, Bhabha et al. 2017). MlaD DNA corresponding to its periplasmic domain (residues 23-183) was chemically synthesised (Genscript) and cloned into pET vector, pET26b (Merckmillipore) to contain a C-terminal 6x His tag. Both MlaC and MlaD plasmids were then used to transform *E. coli* strain BL21(DE3). Expression was performed by growing overnight cultures in lysogeny broth at 37°C to an OD_600_ = 0.6 whereupon protein expression was induced by the addition of IPTG to a final concentration of 1 mM. Protein expression was allowed to continue at 15°C overnight. Cells were harvested by centrifugation (6000 x g, 10 min) and pellets resuspended in 50 mM Tris pH 8, 500 mM NaCl, 10 mM imidazole. Cells were lysed by three passes through an Emulsiflex C3 disruptor (Avestin), then centrifuged (75,000 x g, 45 min) to pellet cell debris. The clarified lysate was passed over a 5 mL Ni-NTA column (GE Healthcare), washed with 5 column volumes of 50 mM Tris pH 8, 500 mM NaCl, 50 mM imidazole and bound protein eluted with 50 mM Tris pH 8, 500 mM NaCl, 500 mM Imidazole. Fractions containing MlaC were further purified on a Superdex 75 gel filtration column (GE Healthcare) equilibrated in 50 mM Tris pH 8, 300 mM NaCl, whilst MlaD containing fractions were additionally purified on a Superdex 200 size exclusion column (GE Healthcare) equilibrated in 50 mM Tris pH 8, 300 mM NaCl.

### Expression and purification of MlaFEDB

MlaFEDB was expressed and purified according to the method of Ekiert *et al*. (2017). Briefly, Rosetta 2 (DE3) cells (Novagen) were transformed with plasmid pBE1196 (Ekiert, Bhabha et al. 2017). For expression, overnight cultures were grown at 37°C with shaking to an OD_600_ of 0.9 then induced by the addition of IPTG to a final concentration of 1 mM for an additional 4 h shaking at 37°C. Cells were harvested by centrifugation (6000 x g, 10 min) and pellets resuspended in 50 mM Tris pH 8, 500 mM NaCl, 10 mM imidazole. Cells were lysed by three passes through an Emulsiflex C3 disruptor (Avestin) before centrifugation (20,000 x g, 30 min) to remove cell debris. The lysate was then further centrifuged (100,000 x g, 30 min) to pellet membranes. Membrane pellets were resuspended and solubilised in 50 mM Tris pH 8, 500 mM NaCl, 10 mM imidazole, 25 mM n-Dodecyl-β-D-Maltopyranoside (DDM) then incubated for 1 h at 4°C with mixing. Insoluble material was pelleted by centrifugation (100,000 x g, 30 min). The supernatant was passed over a 5 mL Ni-NTA column (HIS-Trap – GE Healthcare), washed with 50 mM Tris pH 8, 500 mM NaCl 50 mM imidazole, 0.5 mM DDM and bound protein eluted with 50 mM Tris pH 8, 500 mM NaCl, 500 mM Imidazole, 0.5 mM DDM. Fractions containing MlaFEDB were pooled and concentrated before additional purification on a Superdex 200 size exclusion column (GE Healthcare) equilibrated in 50 mM Tris pH 8, 300 mM NaCl, 0.5 mM DDM.

### Lipid removal

Purified protein was bound to a Ni-NTA affinity column (GE Healthcare) and washed with three washes of 25 mL 50 mM Tris pH 8, 500 mM NaCl, 10 mM imidazole, 25 mM β-octyl glucoside with 1 h static incubation period for each wash, followed by a final wash of 50 mL 50 mM Tris pH 8, 500 mM NaCl, 10 mM imidazole and protein was eluted using 50 mM Tris pH 8, 500 mM NaCl, 250 mM imidazole. Fractions containing protein were pooled, concentrated and then dialysed against 50 mM Tris pH 8, 150 mM NaCl.

### Crystallisation and structure determination of MlaC

Size exclusion fractions containing purified MlaC were concentrated to 20-40 mg·mL^−1^. Vapour diffusion trials were performed via the sitting drop method using the JCSG+ crystal screen (Molecular dimensions). Crystals grew from drops consisting of 2 μl protein plus 2 μl of a reservoir solution consisting of 2.0 M ammonium sulphate, 0.2 M sodium chloride, 0.1 M sodium cacodylate pH 6.5. These were then seeded 1:100, and additional vapour diffusion trials performed using the Morpheus crystal screen (Molecular Dimensions). Crystals grew from drops consisting of 100 nl 1:100 seed solution, 100 nl MlaC (30 mg·mL^−1^), 200 nl reservoir solution consisting of 0.1 M Tris/Bicine pH 8.5, 0.03 M sodium fluoride, 0.03 M sodium bromide, 0.03 M sodium iodide, 20% (v/v) glycerol and 10% (w/v) PEG4000. No additional cryo-protectant was needed. Native diffraction data was collected at Diamond light Source beamline I04, indexed to spacegroup P3_1_21 and processed using XIA2 (Winter 2010). The structure was phased by molecular replacement using PHASER (McCoy, Grosse-Kunstleve et al. 2007) with a search model comprised of the published PL-bound MlaC structure (5uwa). The resulting model was manually rebuilt/adjusted in Coot (Emsley, Lohkamp et al. 2010) and refined using PHENIX (Adams, Afonine et al. 2010) and PDB-REDO (Joosten, Salzemann et al. 2009). The final refined model consists of 2 copies of MlaC within the asymmetric unit, with a final R/Rfree of 19.7/23.6.

### Thin layer chromatography

To analyse PL movement within the Mla system a Ni-NTA affinity column protocol coupled with thin layer chromatography (TLC) was employed. To measure whether MlaC could bind PL directly from the environment, without the need for MlaD or MlaFEDB facilitation, we adjusted both MlaC-lipid and MlaC-apo to 2 mg·mL^−1^ and bound it to a Ni-NTA affinity column equilibrated in 50 mM Tris pH 8, 150 mM NaCl. Small unilamellar vesicles of 1-palmitoyl-2-oleoyl-*sn*-glycero-3-phosphocholine (POPC) or *E. coli* polar lipid extract (Avanti) (4 mg·mL^−1^ in 50 mM Tris pH 8, 150 mM NaCl) were then flowed through the column for 30 min. The column was subsequently washed for 45 min in 50 mM Tris pH 8, 150 mM NaCl, removing any unbound PLs. MlaC was then eluted from the column in 50 mM Tris pH 8, 150 mM NaCl, 250 mM imidazole and collected prior to PL extraction.

To investigate the transport of PLs between MlaC and MlaD equimolar concentrations of each were co-incubated together for 30 min then separated via passage through a Superdex S75 column and fractions containing either MlaC or MlaD collected, analysed by SDS to confirm separation and pooled. Samples were adjusted to 0.5 mg·mL^−1^ in 2 mL and added to 2 mL of methanol and 1 mL chloroform (2:2:1 v/v). Samples were then vortexed continuously for 5 min, incubated for 30 min at 50 °C, and vortexed again for 5 min. The mixture was centrifuged (2000 x *g*, 10 min), and the lower phase extracted and evaporated. Dried PLs were resuspended in 100 μL chloroform and 5 μL loaded onto a Silica TLC plate (Sigma) and run with a 6.5:2:5:1 (chloroform:methanol:acetic acid) solvent. The TLC plate was dried for 30 min, stained with 10 % (w/v) phosphomolybdic acid (PMA) in ethanol, and heated until staining occurred.

To investigate the transport of PLs between MlaC and MlaFEDB, 2 mg of purified MlaFEDB was bound to a 1 mL Ni-NTA affinity column equilibrated in 50 mM Tris pH 8, 150 mM NaCl, 0.5 mM DDM. DDM was then exchanged for 25 mM β-octylglucoside (β-OG) in 50 mM Tris pH 8, 150 mM NaCl. The MlaFEDB bound Ni-NTA affinity column was then incubated for 30 min with 4 mg·mL^−1^ of either POPC or *E. coli* polar lipid resuspended in 50 mM Tris pH 8, 150 mM NaCl, 25 mM β-OG. MlaFEDB was then reconstituted into a PL bilayer environment by flowing POPC or *E. coli* polar lipid small unilamellar vesicles (4 mg·mL^−1^ in 50 mM Tris pH 8, 150 mM NaCl) through the column for 30 min. To remove any unbound PLs MlaFEDB was then thoroughly washed for 45 min in 50 mM Tris pH 8, 150 mM NaCl, with the flow through collected prior to lipid extraction. Either 2 mg MlaC-apo or MlaC-lipid in 50 mM Tris pH 8, 150 mM NaCl was incubated with the bound MlaFEDB on the Ni-NTA affinity column for 30 min. MlaC was then washed off the column in the same buffer and collected for lipid extraction. Bound MlaFEDB was then eluted in 50 mM Tris pH 8, 150 mM NaCl, 250 mM imidazole and collected for lipid extraction.

Samples were adjusted to 0.5 mg·mL^−1^ and 2 mL and added to 2 mL methanol and 1 mL chloroform (2:2:1 v/v). Samples were then vortexed continuously for 5 min, incubated for 30 min at 50°C, and vortexed again for 5 min. The mixture was centrifuged (2000 x g, 10 min), and the lower phase extracted and evaporated. Dried lipids were resuspended in 100 μL chloroform and 5 μL loaded onto a Silica TLC plate (Sigma) and run with a 6.5:2:5:1 (chloroform:methanol:acetic acid) solvent. The TLC plate was dried for 30 min, stained with 10% (w/v) phosphomolybdic acid (PMA) in ethanol, and heated until staining occurred.

### MlaFEDB ATPase activity

An NADH enzyme-linked assay (Norby 1988) adapted for a microplate reader (Kiianitsa, Solinger et al. 2003) was used to determine the rate of ATP hydrolysis by MlaFEDB in both detergent micelles and polar lipid liposomes. All assays were performed in triplicate and at room temperature. For MlaFEDB in detergent micelles, assays were performed in 50 mM Tris pH 8.0, 150 mM NaCl, 5 mM MgCl2, 0.5 mM DDM; whilst for MlaFEDB in liposomes an assay buffer of 50 mM Tris pH 8.0, 150 mM NaCl, 5 mM MgCl2 was used. Using a reaction volume of 75 μl, in 96-well plates (Sigma-Aldrich), 200 mM NADH (Sigma-Aldrich), 20 U·mL^−1^ lactic dehydrogenase (Sigma-Aldrich), 100 U·mL^−1^ pyruvate kinase (Sigma-Aldrich), 0.5 mM phosphoenolpyruvate (Sigma-Aldrich) and different ATP (Sigma-Aldrich) concentrations were added to the appropriate assay buffer. The change in absorbance at 340 nm was then measured using an Anthos Zenyth 340rt (Biochrom) absorbance photometer equipped with ADAP software. Reactions were followed from 10-60 min (allowing for all ATP to be hydrolysed) with measurements taken every 15 s. ATP hydrolysis rates were then determined using maximal 340 nm absorbance readings (with 0 μM ATP), and minimal 340 nm absorbance readings (with 0 μM NADH) from these readings the concentration of NADH at each time-point was calculated, allowing for a linear fit of the reduction in NADH absorbance and conversion to ATP hydrolysis.

### Circular Dichroism

MlaC was diluted to 10 μg·mL^−1^ in 50 mM sodium phosphate pH 7, 300 mM NaCl and circular dichroism performed on a Jasco J-1500 CD spectrometer with a path length of 1 cm at 20°C. A thermal melt was performed with spectra recorded every 5°C up to a temperature of 90°C.

### Analytical ultracentrifugation

Twin channel AUC cells were prepared with 400 μl of protein sample at a concentration of 0.3 mg·mL^−1^ to 1.4 mg·mL^−1^ in one channel and 420 μl of relevant buffer blank in the other. The cells were loaded into an An50Ti rotor in a XL-I analytical ultracentrifuge (Beckman Coulter) and the samples centrifuged at 129,000 x *g* and at a temperature of 20°C until the sample sedimented. The protein samples within the cell were monitored by measuring the absorbance at 280 nm. The data were analysed using the program Sedfit (Schuck 2000) and fit using a continuous c(S) distribution model.

### Nuclear magnetic resonance

NMR experiments were performed at 25°C on an Oxford Instruments 900 MHz magnet equipped with a Bruker Avance III console and TCI 5mm z-PFG cryogenic probe.

Experiments were conducted using ^15^N-labelled MlaC at concentrations between 0.1-0.3 mM in 50 mM sodium phosphate buffer pH 7, 50 mM NaCl and 0.02% (w/v) NaN3 in 90% H_2_O/10% D_2_O. ^15^N labelled MlaC was purified as above but with the exception of substituting lysogeny broth for M9 minimal media containing ^15^NH4Cl as the sole nitrogen source. ^1^H,^15^N HSQC experiments (Kay, Keifer et al. 1992, Palmer Iii, Cavanagh et al. 1992, Schleucher, Schwendinger et al. 1994) were recorded with a spectral width of 7978 Hz and 1024 points in the ^1^H dimension and 1772 Hz and 256 points in the ^15^N dimension in the presence and absence of 2 mg·mL^−1^ *E. coli* polar lipid extract (Avanti). All spectra were processed with NMRPipe (Delaglio, Grzesiek et al. 1995) and analysed using SPARKY (Goddard and Kneller 2004).

### Molecular dynamics simulations

MlaC-apo simulations were conducted using both the MlaC-apo structure determined herein and previously published PL bound MlaC structures from *E. coli* (5uwa), *R. solanacearum* (2qgu) and *P. putida* (4fcz). For all structures, missing terminal residues were added back to the structures prior to simulation using MODELLER, version 9.15 (Fiser and Šali 2003). Prior to simulation the structures were solvated within rhombic dodecahedral boxes, using 100 mM NaCl, while also ensuring system neutrality, and minimised using 1000 steps of the steepest descents method.

Apo simulations initiated from the three different lipid bound MlaC structures were performed using three different force fields: the all-atom AMBER99SBNMR-ILDN (Li and Brüschweiler 2010) and CHARMM36 (Huang and MacKerell 2013) force fields, plus the united-atom GROMOS54A7 force field (Schmid, Eichenberger et al. 2011). Appropriate simulation settings were chosen for each force field. AMBER99SBNMR-ILDN simulations used a 1.0 nm cut-off for both Coulombic and van der Waals interactions. These simulations also used the standard TIP3P water model (Jorgensen, Chandrasekhar et al. 1983). For the CHARMM36 force field, a 1.2 nm cut-off for the Coulombic interactions was applied, while the van der Waals interactions were switched off between 1.0 and 1.2 nm. No long-range dispersion correction was applied. CHARMM36 simulations were performed using the CHARMM TIP3P water model (Durell, Brooks et al. 1994, Neria, Fischer et al. 1996). GROMOS 54A7 simulations used a 1.4 nm cut-off for both Coulombic and van de Waals interactions and the SPC water model (Berendsen, Postma et al. 1981). Unlike the all-atom force field simulations, which used a simulation time step of 2 fs, the GROMOS 54A7 force field simulations used virtual interaction sites for the hydrogen atoms allowing a 4 fs time step to be applied (Feenstra, Hess et al. 1999). For all these simulations: long-range interactions were treated using the Particle Mesh Ewald method (Essmann, Perera et al. 1995); a simulation temperature of 310 K and a pressure of 1 bar were maintained using Nose-Hoover (Nosé 1984, Hoover 1985) and Parrinello-Rahman (Parrinello and Rahman 1981, Nosé and Klein 1983) schemes with coupling constants of 2.0 and 5.0 ps, respectively; for each protein and force field combination, five independent simulations utilising different starting velocities were performed for 500 ns each, following an initial 10 ns period in which the positions of the protein atoms were restrained. These simulations were all performed using GROMACS, version 5.0.6 (Abraham, Murtola et al. 2015).

Simulations initiated from the MlaC-apo structure were performed using the AMBER99SBNMR-ILDN force field and followed the same settings as described above. Two standard simulations were performed for 500 ns, each using different starting velocities. Simulations were also performed using both elevated temperature and scaled down interactions, as per the higher replica levels of temperature and Hamiltonian replica exchange methods (Sugita and Okamoto 1999, Bussi 2014), in an attempt to accelerate the sampling within these simulations. The elevated temperature simulations were performed at a temperature of 400 K, while the interactions were scaled down by a factor of 0.7. We note here that applying a higher temperature of 500 K or a scaling factor of 0.5 led to the protein unfolding during test simulations. As per the standard MlaC-apo simulations, two independent simulations were performed for each setting using different starting velocities. Scaling of the interactions was performed using the partial_tempering script of the PLUMED code (Tribello, Bonomi et al. 2014). Apart from the higher temperatures and scaled interactions, these simulations were performed using the same methods described for the standard simulations. Simulations of the MlaC-apo structure were all performed using GROMACS version 5.1.4 (Abraham, Murtola et al. 2015). Analysis of the simulations was performed using the GROMACS tools, the trj_cavity program (Paramo, East et al. 2014) and VMD (Humphrey, Dalke et al. 1996).

### Functionalisation of gold surfaces

Quartz sensors with gold surface coating were cleaned by UV-ozone treatment for 10 min then immersed in a 5:1:1 solution of H_2_O:H_2_O_2_ (25%):NH4OH (30%) heated to 75 °C for 5 min. Sensors were thoroughly rinsed in ultra-pure water, dried and subject to a final UV-ozone treatment for 10 min. The gold surfaces were then functionalized based on the method of Giess (Giess, Friedrich et al. 2004). Briefly, surfaces were immersed in a solution of 2 mg·mL^−1^ dithiobis (*N*-succinimidyl propionate) in dry DMSO for 30 min, then rinsed with dry DMSO, ultra-pure water, and ethanol and dried in a stream of nitrogen. The surfaces were then immersed for 2 h in a solution of 150 mM *N*-(5-amino-1-carboxypentyl) iminodiacetic acid (ANTA) buffered to pH 9.8 with 0.5 M K2CO3, washed with ultra-pure water and then incubated for 30 min in 40 mM CuSO4 in 50 mM sodium acetate buffer. As a final step, the surfaces were washed with ultra-pure water and dried in a nitrogen stream prior to use.

### Quartz crystal microbalance with dissipation monitoring (QCM-D)

QCM-D was performed using a Q-sense Analyzer QCM-D system (Gothenburg, Sweden). Sensors were mounted into a temperature controlled flow cell at 20°C attached to a calibrated peristaltic pump and filled with ultra-pure H_2_O. A flow rate of 0.1 mLomin^−1^ was used throughout unless otherwise stated. A baseline was acquired for 5 min in H_2_O to allow for temperature equilibration before the measurement was started. Buffer was exchanged for DDM buffer (50 mM Tris pH 8, 150 mM NaCl, 0.5 mM DDM) for 10 min. MlaFEDB with a 6 x His tag at the N-terminal of MlaF was diluted to 0.1 mg·mL^−1^ in the same buffer and injected into the cell to bind to the Cu^2+^ surface. Once Δ*f* reached −50 Hz, excess MlaFEDB was removed by flowing DDM buffer over the surface for 7 min. DDM was exchanged for β-octylglucoside (β-OG) (25 mM in 50 mM Tris pH 8, 150 mM NaCl) for 15 min to ensure thorough detergent exchange. Mixed POPC and β-OG micelles (0.2 mg·mL^−1^ POPC, 25 mM β-OG, 50 mM Tris pH 8, 150 mM NaCl) were injected for 15 min to allow for detergent to exchange for PLs. POPC small unilamellar vesicles (0.2 mg·mL^−1^ in 50 mM Tris pH 8, 150 mM NaCl) were then flowed for 10 min to deposit a supported bilayer. The surfaces were thoroughly washed for 20 min in 50 mM Tris pH 8, 150 mM NaCl to ensure excess PL had been removed. MlaC was diluted to 0.1 mg·mL^−1^ in 50 mM Tris pH 8, 150 mM NaCl in the presence or absence of 0.5 mM ATP, 5 mM MgCl2 for ATP dependence experiments. MlaC was injected into the cell at a flow rate of 0.1 ml·min^−1^. Excess MlaC was removed by washing the surfaces in 50 mM Tris pH 8, 150 mM NaCl. This allowed Δ*f* in response to MlaC to be obtained.

### Attenuated total reflection Fourier transform infrared spectroscopy (ATR-FTIR)

ATR-FTIR spectra were collected using a Thermo-Nicolet iS50 instrument fitted with an ATR flow cell accessory (Specac) attached to a syringe pump and a water cooling loop connected to a temperature controlled water bath, a cryo-cooled mercury cadmium telluride detector and a dry-air purge operating at a flow rate of 40 L·min^−1^ in order to minimise absorbance from residual water vapour in the beam path. All spectra were obtained at 20 °C, with a resolution of 4 cm^−1^ and 128 interferograms collected for each spectra. Fourier self-deconvolution was performed automatically by the data acquisition software (OMNIC 9 - Thermo Fisher Scientific) with a constant bandwidth applied across all spectra.

Single crystal silicon ATR substrates were cleaned by immersion in 2% (w/v) SDS for 30 min before rinsing extensively with ultrapure water and drying under a stream of nitrogen. Substrates were then UV-ozone cleaned for 10 min, washed with ultrapure water and then UV-ozone cleaned a final time for 10 min. The substrate was mounted in the flow cell dry and the volume filled with buffer A (50 mM Tris, 150 mM NaCl, pD 8.0 in D_2_O) for background collection. For experiments investigating the effect of ATP, all buffers were supplemented with 0.5 mM ATP, 5 mM MgCl2. Proteoliposomes for deposition were prepared by mixing POPC and MlaFEDB in a 3:1 mass ratio in the buffer B (50 mM Tris, 150 mM NaCl, 25 mM β-OG, pD 8.0 in D_2_O) to a final concentration of 0.3 mg/mL MlaFEDB and 0.1 mg/mL POPC. This mixture was then diluted 10-fold in buffer A to lower the detergent concentration below the critical micelle concentration, and incubated at 6 °C for 30 min. This resulted in spontaneous self-assembly of proteoliposomes of approximately 100 nm diameter, as confirmed by dynamic light scattering. 2 mL POPC/MlaFEDB proteoliposomes were then manually injected over the Si substrate and incubated for 15 min to allow for MlaFEDB-containing POPC bilayer deposition. The proteoliposome suspension was diluted a further 10-fold in buffer A and 2 mL injected into the flow cell to allow for β-OG to exchange out of the deposited bilayer. Finally, the POPC/MlaFEDB bilayer was washed with 5 mL buffer A at a flow rate of 0.5 mL·min^−1^ and spectra collected.

MlaC was introduced into the cell at a concentration of 10 μg·mL^−1^ in buffer A under a constant flow of 0.1 mL·min^−1^ for 4 h. Spectra were continuously collected every 80 s throughout. After four hours, the flow cell was washed with 5 mL buffer A at a flow rate of 0.5 mL·min^−1^ and a final spectrum collected.

All spectra were corrected for removal of water vapour by scaling and subtracting spectra collected prior to deposition of the POPC/MlaFEDB bilayer from the bilayer spectra. No further processing was performed. Peak integrations were performed over the aliphatic C-H stretch region from 2990 cm^−1^ – 2810 cm^−1^. This includes contributions from the symmetric and asymmetric C-H2 stretching vibrations and the C-H3 stretching vibrations arising from the PL tails as a qualitative comparison of the extent of PL removal from the bilayer.

## Results

### Detergent treatment of MlaC can produce a phospholipid free variant

The current model of Mla-mediated PL transport implies the presence of a periplasmic component (MlaC) that is able to ferry PL from the OM to the IM (Figure 1). For this model to be correct MlaC must be able to adopt two forms, PL bound and PL free. All previous studies of MlaC (and its homologues) have isolated the protein in its PL-bound form (Thong, Ercan et al. 2016, Ekiert, Bhabha et al. 2017). In order to investigate the directionality of PL transport, we required the isolation of both species. Using *E. coli* pBE1203; (Ekiert, Bhabha et al. 2017)) we overexpressed and purified MlaC by published methods (Ekiert, Bhabha et al. 2017) (Figure S1). Thin layer chromatography (TLC) analysis of MlaC extracts revealed MlaC co-purified with PL with a particular preference for phosphatidyl glycerol, as observed previously (Thong, Ercan et al. 2016). However, unlike previous observations we also noted the presence of cardiolipin (Figure S2a). Throughout this study we term this species MlaC-lipid. In order to produce protein free of bound PL (MlaC-apo), we washed the protein extensively using buffer containing the detergent β-octyl glucoside. TLC analyses confirmed that PL had been completely removed from the sample (Figure S2A) whilst circular dichroism spectroscopy indicated that secondary structure content of MlaC-apo was indistinguishable from MlaC-lipid (Figure S2B). However, a 3°C decrease in thermal stability from 72.5°C to 69.5°C was observed (Figure S2D) suggesting that PL binding stabilises the protein structure. Analytical ultracentrifugation showed no significant changes in oligomeric state (Figure S2C), with Svedburg values of 1.92 and 1.97 S for MlaC-apo and MlaC-lipid respectively. The fact that the overall structure of MlaC remained unperturbed by PL removal was confirmed by NMR spectroscopy. Both MlaC-lipid and MlaC-apo showed a single folded species as characterised by the expected number of peaks observed and good dispersion of signals (Figure S2E). However clear differences in peak positions were noted across a number of residues suggesting a conformational change occurred during PL addition/removal.

### The phospholipid binding cavity of MlaC is controlled via a β-sheet hinged lid

To determine the structural changes that occur in MlaC upon loss of PL, we determined the crystal structure of MlaC-apo at a resolution of 2.25 Å (Table S1). Within the asymmetric unit two molecules of MlaC were observed, one in a conformation similar to the published lipid-bound MlaC structure (5uwa) but lacking density associated with PL, which we have termed MlaC-open and another in a previously undescribed closed conformation (MlaC-apo) (Figure 2A & B). Both structures maintained secondary structural elements observed for the published PL bound crystal structure (5uwa) with a mixed α/β-fold consisting of a highly twisted four-stranded β-sheet and a bundle of seven α helices. The interaction between the two molecules was relatively small ~194 Å^2^ (MSMS) and the AUC analyses indicated that both MlaC-apo and MlaC-lipid are monomeric. These data suggest the observed interaction surface is not physiological.

**Figure 2.**
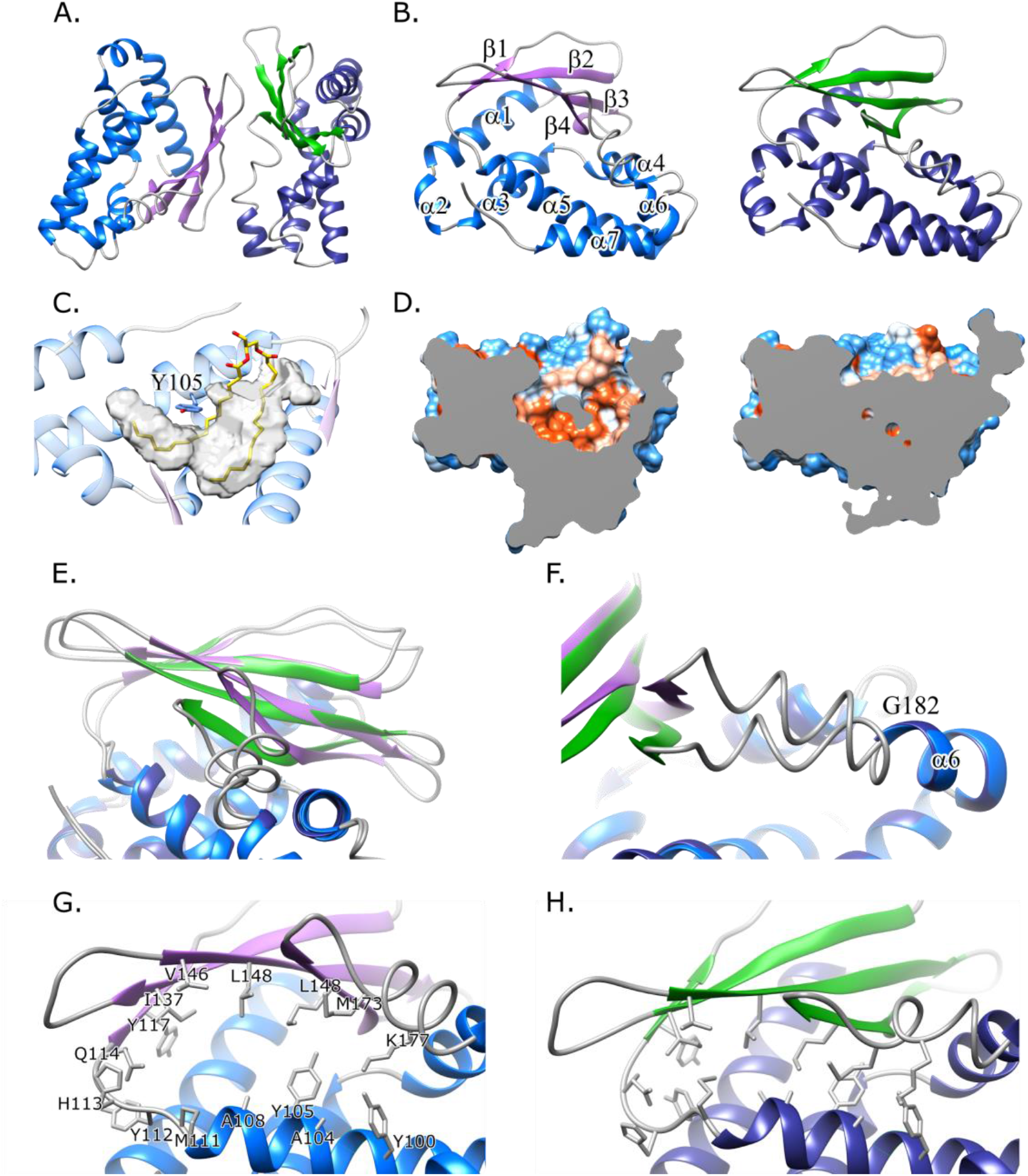
The MlaC-apo structure. A) Cartoon representation of the two molecules of MlaC observed in the asymmetric unit. The MlaC-open conformation is shown coloured light blue (helix) and pink (sheet) and the closed MlaC-apo conformation coloured dark blue (helix) and green (sheet). B) The individual open and apo conformations of MlaC highlighting the numbering of secondary structural elements. C) Stripped away structure showing the MlaC PL binding pocket observed in the MlaC open conformation overlaid with the position of the PL observed in crystal structure 5uwa highlighting the position of Y105 bisecting the pocket. PL binding pocket calculated using 3vee (Voss and Gerstein 2010). D) Cut away surface representation coloured according to hydrophobicity (Blue hydrophobic, white – neutral, red – hydrophilic) of both the MlaC open (left) and apo (right) conformations showing the closing of PL binding pocket. E) Observed shift of the β-sheet between the open and apo conformations with a lateral movement of 2.5A and 15° shift in angle occurring. F) Corresponding shift of part of helix α6 pivoting from Glyl82 resulting in an angle change of 17.9°. Zoom in view of the residues surrounding the binding pocket in the open (G) and apo (H) conformations showing little conformational change in the residues involved.

The open conformation gives an RMSD for Cα of 0.77 Å and 0.63 Å against the two MlaC molecules observed in the asymmetric unit of the PL bound crystal structure (5uwa). Analysis of the cavity observed in MlaC-open shows it is very similar to that of 5uwa and overlaps closely with the position of the bound PL in 5uwa, with Tyr105 also shown to bisect the cavity (Figure 2C). On further analysis of the structure, however, it was evident that this open conformation was likely the result of crystal packing between the two conformations within the asymmetric unit; with the β-sheet on the pocket side of the pivot in the open conformation making contact with MlaC-apo, pulling the pocket open.

Unlike MlaC-open, MlaC-apo shows the absence of a pocket (Figure 2D). Comparison between the apo and open conformations indicate that the majority of the protein undergoes only minor conformational changes (0.66Å over α helices 1-5 & 7; residues 23-112,183-208) between the open and apo conformations. However, large changes are observed across the β-sheet, in particular a ~2.5 Å translation of the entire β-sheet and a pivot of approximately 15° (Figure 2E). To accommodate this movement a further pivot is manifest at Gly182 within helix α6 resulting in an angular shift of part of this helix by 17.9° (Defined across residues 169-182) (Figure 2F). These movements together allow the residues at the mouth of the cavity to come together, which results in its closing (Figure 2G & H).

To corroborate the MlaC-apo structure a series of fifteen independent molecular dynamics simulations, initiated from the PL bound crystal structure with the PL removed (5uwa) were performed. Within all simulations, irrespective of the force field used, there was a rapid collapse of the PL binding cavity within the first 25 ns of the simulations, and movements of the protein towards the observed MlaC-apo structure (Figure S3 and Table S2). Additional simulations performed of the MlaC-apo structure demonstrate the stability of the protein in this apo conformation (Figure S4A and B) and further show the close similarity of the MlaC-apo structure to the structures identified in the simulations with the bound PLs removed (Figure S4C). Furthermore, a further series of PL removed simulations performed on MlaC structures determined from *Ralstonia solanacearum* (2qgu) and *Pseudomonas putida* (4fcz) demonstrate a very similar closure of the PL binding pocket upon PL removal (Figure S5). In particular there are pivoting movements of the β-sheet and movements of the C-terminal end of helix α5 which together dominate the changes observed upon pocket closure. This rapid closure is also in agreement with previous simulations of the *R. solanacearum* structure (Huang, Miao et al. 2016).

### MlaC cannot bind free phospholipid

We next investigated whether MlaC-apo is able to bind passively to free PL. To this end, we incubated hexahistidine-tagged MlaC-apo with buffer containing *E. coli* PLs and removed the unbound PLs by purifying MlaC using Ni-NTA resin. Analysis of the Ni-NTA eluate by TLC indicated that MlaC-apo did not bind free PL (Figure 3A). To more accurately confirm this we probed PL binding using NMR. By comparing spectra taken of both ^15^N-MlaC-lipid and ^15^N-MlaC-apo we were able to monitor any changes in binding on incubation of MlaC-apo with excess *E. coli* polar lipids (Figure 3B & S6). Following co-incubation for 18 h at room temperature minimal PL loading occurred, as evidenced by chemical shifts predominantly remaining in the apo position. These data suggest that loading is either an extremely slow process or requires interaction with another Mla component. Consistent with these results molecular dynamics simulations of MlaC-apo performed to advance the conformational sampling demonstrates that once the structure is closed and in its apo conformation it does not readily sample the open conformation, even at higher temperatures or in simulations with scaled-down interactions (Figure S7). These studies support the observation that the MlaC-open conformation is the result of crystal packing.

**Figure 3.**
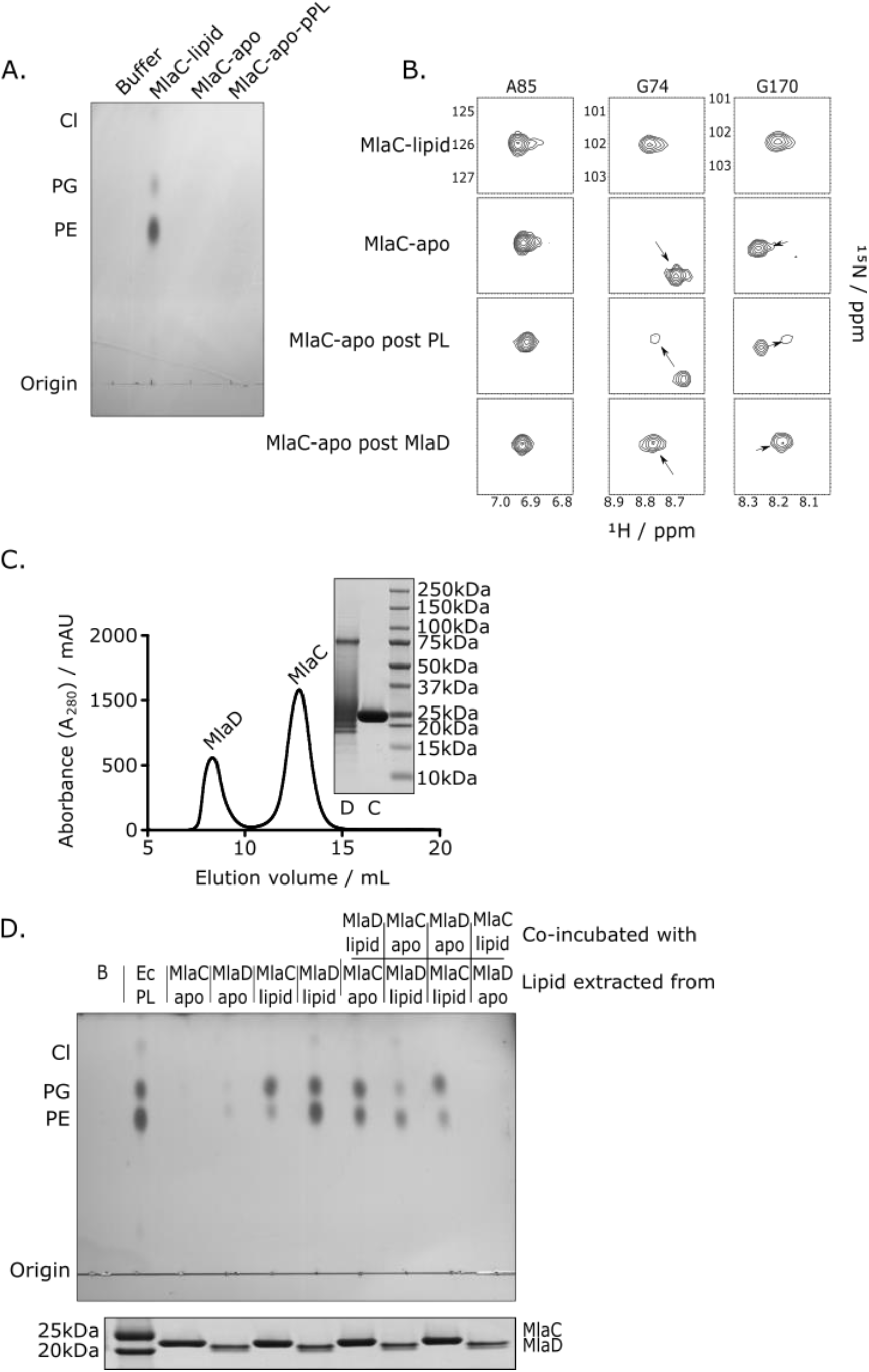
MlaC requires MlaD for phospholipid loading. A) TLC of MlaC purified from the cell showing the presence of bound PL (MlaC-lipid); following washing with detergent leading to PL removal and the formation of MlaC-apo; and MlaC apo following incubation with *E. coli* polar lipids (MlaC-apo-pPL) showing no addition of lipid to MlaC. B) Excerpts from ^1^H^15^N-HSQC spectra of 0.2mM ^15^N-labelled MlaC showing the H-N resonance positions for A85, G74 and G170 in the presence of PL (MlaC-lipid), the absence of PL (MlaC-apo), following 18 h incubation of MlaC-apo with 2mg·mL^−1^ *E. coli* polar lipids (MlaC-apo post PL), and following 30 min incubation of MlaC-apo with MlaD-lipid and subsequent SEC purification (MlaC-apo post MlaD). C) Size exclusion chromatogram of the purification of MlaC and MlaD following co-incubation to allow PL transfer to occur. Inset shows the unboiled SDS-PAGE of the components following separation with MlaD shown forming characteristic oligomers (Thong, Ercan et al. 2016, Ekiert, Bhabha et al. 2017). D) TLC showing the movement of PL between MlaC and MlaD. Positions of phosphatidyl glycerol (PG), phosphatidyl ethanolamine (PE) and cardiolipin (Cl) are shown. Below is shown an extract of an SDS-PAGE showing the separation of MlaC and MlaD for each sample.

### MlaC accepts phospholipid from MlaD

Malinverni & Silhavy (2009) suggest that MlaC passes PLs to the MlaFEDB complex (Figure 1). Because the MlaD component of this complex has been shown to interact with MlaC (Ekiert, Bhabha et al. 2017) we sought to investigate PL movement between these two components. Thus we overexpressed and purified a construct of MlaD consisting of its periplasmic domain lacking its N-terminal membrane anchor (residues 29-183) with a C-terminal 6 x His tag (Figure S8). Like MlaC, MlaD copurified with PLs (MlaD-lipid) (Figure 3D). We generated PL free MlaD (MlaD-apo) using the same method we described for MlaC-apo (Figure 3D). Next we probed PL transfer between MlaC and MlaD by co-incubation of apo and PL forms. After co-incubation each protein species was separated by size exclusion chromatography and their PL content analysed by TLC (Figure 3C). Unexpectedly, no lipid transfer was observed between MlaC-lipid and MlaD-apo. In contrast, lipid transfer from MlaD to MlaC was clearly observed when MlaC-apo was co-incubated MlaD-lpid (Figure 3D) as evidenced by PL association with MlaC and diminished levels of PL associated with MlaD. To confirm the transfer event yielded a structure consistent with MlaC-lipid we again utilised NMR. Analysis of ^15^N-MlaC-apo following incubation with MlaD-lipid and subsequent purification showed that MlaC adopted the MlaC-lipid conformation (Figure 3B & S6A). However incubation of ^15^N-MlaC-lipid with MlaD-apo showed no change in chemical shift confirming the TLC results. Owing to the passivity of these interactions (i.e. in the absence of an energy source) the clear unidirectional movement of PL to MlaC suggests that it has a far higher binding affinity for PL than MlaD.

### MlaFEDB transfers phospholipid from the inner membrane to MlaC

Given that MlaC has a higher affinity for lipid than MlaD, we hypothesised that the full MlaFEDB complex may increase the affinity of MlaD for PL or promote uptake of PL from MlaC. MlaFEDB has been established as an ATP-binding cassette (ABC) family transporter (Thong, Ercan et al. 2016). Using a similar construct to those previously published (Thong, Ercan et al. 2016, Ekiert, Bhabha et al. 2017) but with a Histidine affinity tag present at the N-terminal of MlaF, we purified the complex in detergent micelles (Figure 4A) and were able to show intrinsic ATPase activity (Figure 4B) in line with others (Thong, Ercan et al. 2016). By binding MlaFEDB to metal chelate resin, we were able to stably reconstitute the complex back in to PL. This allowed us to perform interaction studies by flowing MlaC over the complex (in the presence of ATP), which could then be analysed for the presence of bound PLs (Figure 4C & D). The results corroborated the observations for MlaD and again showed the movement of PLs from MlaFEDB to MlaC-apo. Movement of PL from MlaC-lipid to MlaFEDB was not observed. To confirm that binding to the resin resulted in no adverse effects on ATPase functionality we measured ATPase activity whilst bound to resin beads (Figure S9). Only a small decrease was noted, suggesting tethering had no adverse effects on function.

**Figure 4.**
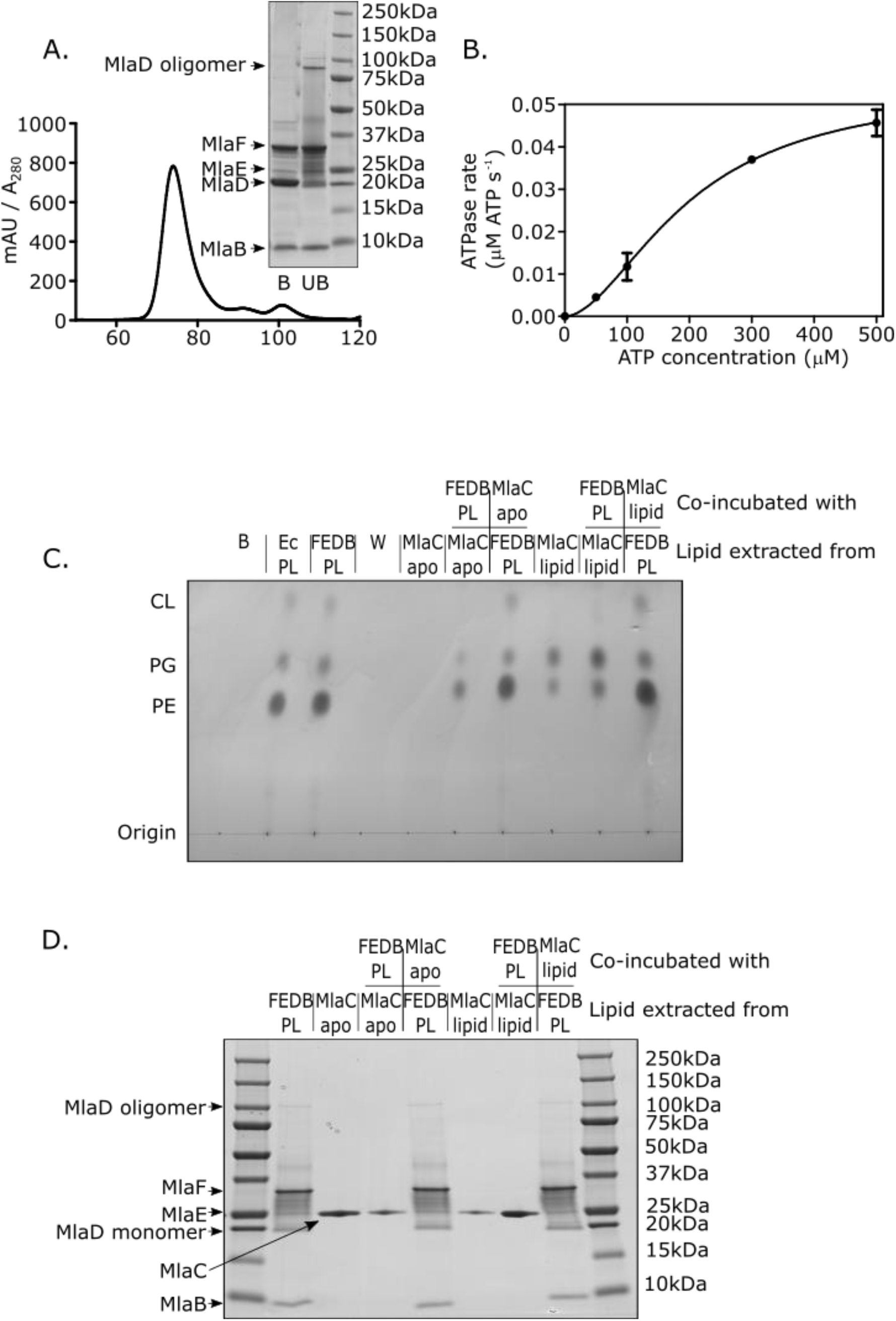
MlaFEDB is active and exports inner membrane phospholipids to MlaC. A) Size exclusion chromatogram of detergent solubilised MlaFEDB yielding a single complex. Inset showing SDS-PAGE demonstrating that MlaF, MlaE, MlaD and MlaB form a complex, with characteristic oligomers of MlaD shown in the unboiled (UB) sample which are lost in the boiled (B) sample. B) Enzyme coupled ATPase assay of MlaFEDB (0.1μM) performed in detergent micelles (0.05% DDM). Average ATP hydrolysis rates (obtained from triplicate experiments) were plotted against ATP concentrations and fitted against an expanded Michaelis-Menten equation that includes a term for the Hill coefficient (n). V_max_ = 0.05 μM ATP s^−1^, k_*cat*_ = 0.5 ± 0.1 s^−1^, K_m_ = 200.0 ± 44.6 μM, n = 1.81 ± 0.35. C) TLC showing the movement of PL between MlaFEDB and MlaC in the presence of 500 μM ATP. B – Buffer, Ec PL – *E. coli* Polar lipids; FEDB PL – MlaFEDB reconstituted in to *E. coli* polar lipid liposomes, W – Wash following binding of MlaFEDB in *E. coli* polar lipid liposomes to Ni-NTA column. D) SDS-PAGE of the samples shown in C) confirming separation of the various species following incubation.

To further address and confirm transfer directionality and provide a method to investigate rate changes we utilised the technique of quartz crystal microbalance with dissipation monitoring (QCM-D). QCM-D is a surface sensitive technique utilising the piezoelectric effect for real-time measurements of surface interactions. The change in the resonant frequency (Δ*f*) is inversely proportional to the amount of mass deposited at the surface (Δ*m*), as given by the Sauerbrey equation (Sauerbrey 1959):

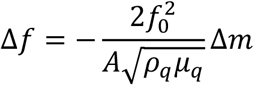

Where *f*_0_ is the resonant frequency, *A* is the piezoelectrically excited area of the crystal, *ρ_q_* and *μ_q_* are the density and shear modulus of the quartz crystal, respectively.

Using the approach of Giess *et al* a protein tethered PL bilayer was generated (Giess, Friedrich et al. 2004). This required attaching MlaFEDB to an engineered gold surface modified with Cu^2+^-NTA followed by reconstitution of a PL bilayer around the complex (Figure 5A). Copper was chosen over Nickel to provide a stronger interaction with the His-tag, again this did not overly affect activity (Figure S9). Using the same construct of MlaFEDB with a 6xHis tag on the N-terminal of MlaF, which we had already shown was active, allowed for correct orientation on the surface. Addition of detergent solubilised MlaFEDB resulted in a decrease in frequency corresponding to a mass uptake (Figure 5B-1). The surface was then washed (Figure 5B-2), exchanged in to a β-octyl glucoside detergent buffer that more suited vesicle deposition (Figure 5B-3) but which did not inhibit function (Figure S10). A supported bilayer was formed by first an addition of mixed detergent/POPC (Figure 5B-4) and finally POPC alone leading to a characteristic initial adsorption and subsequent rupture of vesicles indicative of bilayer deposition at the surface (Richter, Mukhopadhyay et al. 2003) (Figure 5B-5) and final washing to remove excess liposomes (Figure 5B-6). Because the Mla pathway does not appear to show head group preference (Ekiert, Bhabha et al. 2017), POPC was chosen due to its uniform and well characterised properties and known surface deposition (Shen, Leyton et al. 2014). Although not a constituent of *E. coli* membrane, MlaFEDB functionality was not impaired by the presence of POPC (Figure S11A & B). We also confirmed that MlaC is not able to passively take up free POPC (Figure S11C). We then sought to see if the addition of MlaC-apo or MlaC-lipid, in the presence of ATP, would either lead to deposition or removal of PLs from the surface through MlaFEDB, after first confirming that no interaction takes place with the surface in the absence of MlaFEDB (Figure 5C). Addition of MlaC-lipid resulted in no persistent changes in mass (Figure 5B-7 & 5B-8, Black) suggesting no addition of PL to the bilayer occurred. However, on addition of MlaC-apo clear increases in frequency were observed (Figure 5B-7 & 5B-8, Red), consistent with mass loss from the surface, and taken with the TLC is consistent with the removal of PL from the bilayer via MlaFEDB.

**Figure 5.**
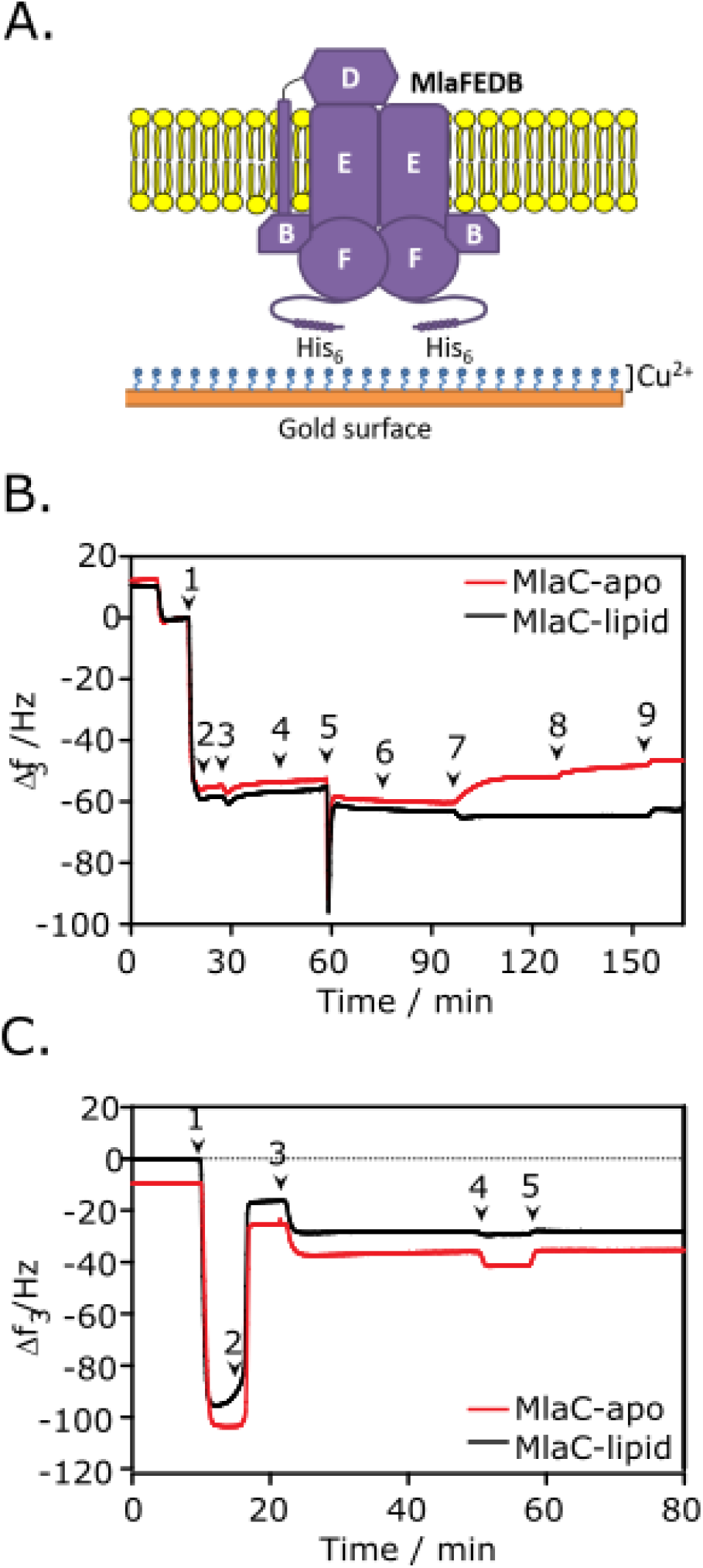
QCMD further suggests MlaFEDB exports inner membrane phospholipids to MlaC. A) Schematic showing how MlaFEDB associates via its 6 His tag with the Cu^2+^ functionalized gold surface of the QCM-D sensor and is reconstituted within a membrane bilayer. B) QCM-D measurements performed in the presence of 500μM ATP showing the frequency response, indicative of mass changes to [1] the attachment of MlaFEDB to the gold surface, [2] washing off excess MlaFEDB, [3] exchange of DDM detergent for β-OG detergent to facilitate PL deposition, [4] exchange of β-OG detergent with a mixture of β-OG and POPC, [5] POPC vesicle deposition and popping, [6] washing off excess POPC vesicles with Tris buffer, [7] addition of MlaC either apo (red) or lipid bound (black) followed by flow stop, [8] repeat injection of MlaC followed by flow stop, [9] buffer wash. C) QCM-D measurements showing the frequency response, indicative of mass changes to [1] POPC PL deposition [2] washing with water to promote vesicle popping, [3] washing with buffer, [4] addition of purified MlaC either apo (red) or lipid bound (black), [5] final wash with buffer to remove any bound MlaC.

To confirm that the mass loss observed from the surface was due to PL and not protein we employed the use of attenuated total reflection (ATR) fourier transform infra red (FTIR) spectroscopy (Figure 6). Briefly, FTIR spectroscopy provides a qualitative analysis of the chemical bonds present within a sample by their characteristic bond vibrational frequencies.

**Figure 6.**
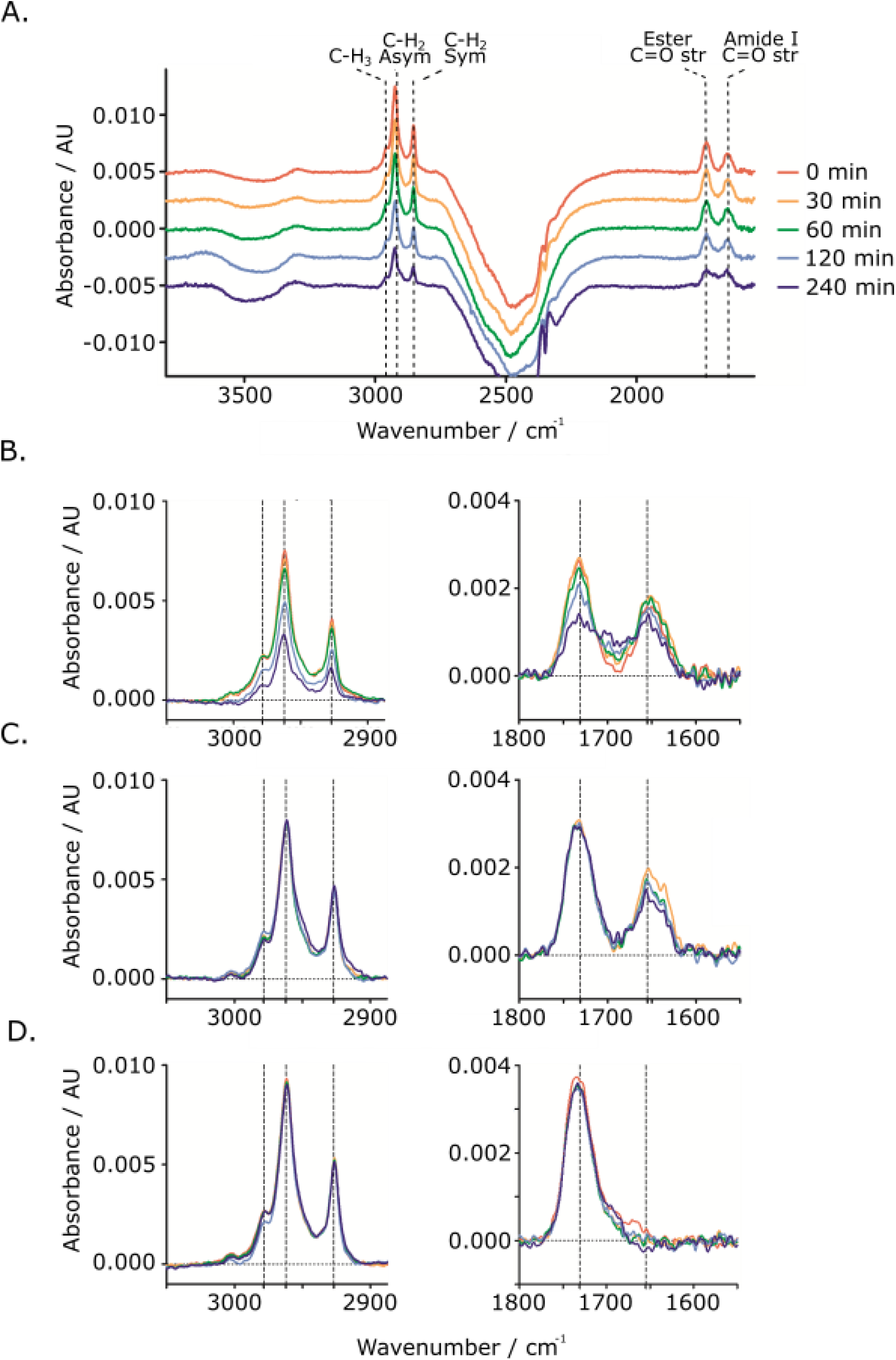
FTIR confirms MlaFEDB extracts phospholipid from the membrane and passes it to MlaC-apo. A) ATR-FTIR spectra of a Si-adsorbed supported POPC bilayer containing reconstituted MlaFEDB, offset with respect to each other for clarity. Spectra were collected before interaction with MlaC-apo (red spectra) and then at selected time points during interaction with MlaC-apo (orange through indigo spectra). Dashed vertical lines highlight absorbances corresponding to stretching vibrations of symmetric and asymmetric C-H2 and C-H3 groups, present primarily within the POPC tails, the ester C=O stretch, from the glycerol-ester bond of the PL tails, and the amide I band from the HNC=O bonds in the protein backbone. The strong negative absorbance band at 2500 cm^−1^ is due to fewer O-D bonds from the D_2_O buffer within the penetration depth of the IR effervescent wave after bilayer deposition relative to background spectra collected of bare Si. B, C) Excerpts from ATR-FTIR spectra showing the absorbance of aliphatic C-Hx bonds (left panel) and carbonyl bonds (right panel) during the interaction of Si-adsorbed MlaFEDB/POPC bilayers with B) MlaC-apo or C) MlaC-lipid. D) Excerpts of ATR-FTIR spectra of a Si-adsorbed POPC bilayer taken at selected time points whilst in the presence of MlaC-apo. All spectra displayed are taken at timepoints as indicated in Figure 6A and collected in the presence of 500 μM ATP throughout.

Whilst ATR-FTIR spectroscopy allows an IR spectrum to be recorded from the reflection from the surface of a substrate and provides information on the chemical composition of that surface. Unlike QCM-D we were not able to utilise gold to orientate the protein as the IR beam could not penetrate sufficiently. However, we found we were able to simply deposit active MlaFEDB proteoliposomes on the silicon oxide surface of a single-crystal Si ATR substrate, which we confirmed by QCM-D resulting in similar results to those shown in Figure 5. A clear distinction between PL and protein could be achieved by the presence of the unique protein amide-I C=O vibration at ~1650 cm^−1^ and the corresponding PL molecular vibrations of C=O ester, CH3, CH2 asymmetric and CH2 symmetric stretching vibrations at 1731, 2855, 2924 and 2958cm^−1^, respectively (Figure 6A). On passage of MlaC-apo over the supported bilayer surface, clear decreases in vibrations associated with PL were observed, with no concomitant drop in protein signal (Figure 6B), such loss was not observed in the presence of MlaC-lipid (Figure 6C), nor for MlaC-apo in the presence of POPC alone (Figure 6D) confirming that the mass loss from the surface observed in QCM-D was the result of PL loss only. By measuring changes in peak area, over the 4 h interval approximately 36% of PL was removed from the surface, but it must be noted that in this system MlaFEDB orientation was not defined, it could adopt one of two orientations, either with MlaD exposed on the surface or with MlaF/B exposed, thus likely leading to an underestimation of the percentage of PL loss relative to the correctly orientated, tethered MlaFEDB bilayer system. Furthermore, no accurate information regarding the concentration of MlaFEDB on the surface could be determined. To identify whether MlaFEDB functions in an enzymatic fashion, e.g. can undergo multiple rounds of PL extraction, we utilised QCM-D with continuous passage of MlaC-apo over the surface (Figure 7B). Using the Sauerbrey equation (Sauerbrey 1959), and based on mass loss being due solely to PL removal and assuming no changes in viscoelasticity, over the experimental time course (40 min) and concentration of MlaC-apo used (0.01 mg·mL^−1^), we estimated that roughly 90 molecules of PL were extracted per MlaFEDB complex, strongly suggesting that MlaFEDB functions in an enzymatic fashion.

**Figure 7.**
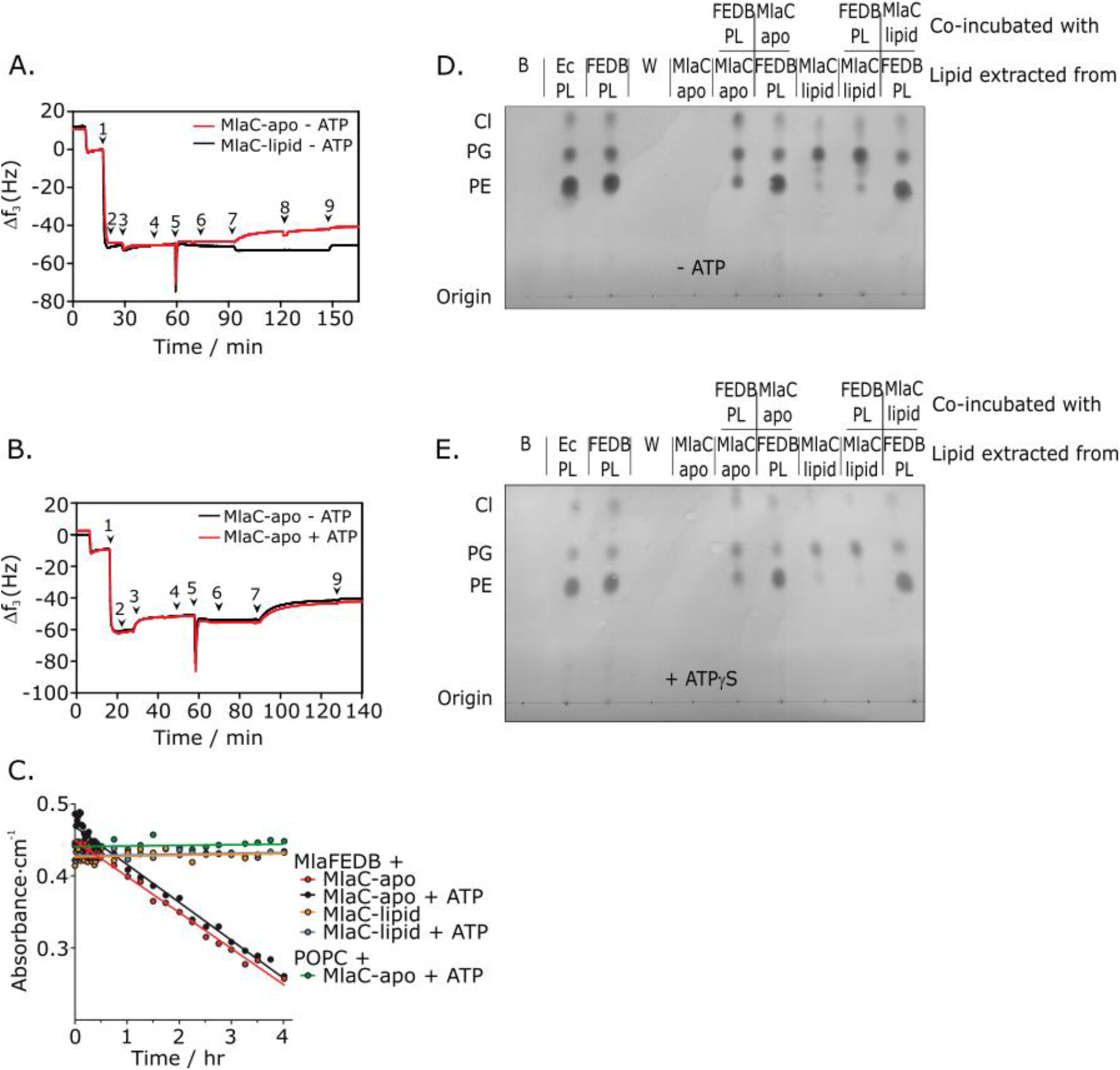
MlaFEDB is able to export phospholipids in an ATPase independent manner. A) QCM-D measurements showing the frequency response observed as in Figure 5B but in the absence of ATP. B) Side-by-side comparison of QCM-D profiles in the presence (Red) and absence (Black) of ATP with numbering according to Figure 5B. C) Changes to ATR-FTIR integral values for aliphatic C-H_x_ peaks (2900 – 2810 cm^−1^) of Si-adsorbed MlaFEDB/POPC supported bilayers upon introduction of MlaC (apo/lipid) in the presence or absence of ATP. D) TLC showing the movement of PL between MlaFEDB and MlaC in the absence of ATP labelled according to Figure 4C. E) TLC showing the movement of lipid between MlaFEDB and MlaC in the presence of 500 μM ATPγS, labelled according to Figure 4C.

### ATPase activity increases in the presence of MlaC-apo

Thong *et al*. (2016) highlighted the ability of MlaD to moderate the ATPase activity of the MlaFEDB complex, and suggested that MlaC might play a regulatory role, potentially increasing the MlaFEDB ATPase activity via direct interactions at the PL bilayer. To establish the role of the PL environment and probe the ability of MlaC to influence MlaFEDB ATPase activity, MlaFEDB was introduced into *E. coli* polar lipid liposomes and subsequently incubated with either MlaC-apo or MlaC-lipid, the ATP hydrolytic activity of MlaFEDB was then assessed through an enzyme-coupled ATPase assay (Norby 1988). Following the introduction of MlaFEDB into polar lipid liposomes, ATPase activity was dramatically diminished with the rate at 500 μM ATP decreasing from 0.05 μM ATP s^−1^ (MlaFEDB in detergent micelles) to 0.009 μM ATP s^−1^ (Figure 8). This is not surprising, if ATP has a role in PL transport, when removed from detergent and in the membrane bilayer MlaFEDB loses the ability to load or off-load PLs as the aqueous environment no longer harbours water soluble lipid like detergents which can bind or be freely released from MlaFEDB. Instead they are replaced by hydrophobic PLs that preferentially form stable liposomes. Thus for PL transport to occur a donor or acceptor must be present, e.g. MlaC. Following incubation with MlaC-lipid, the rate was unaffected. However, upon addition of MlaC-apo, the ATPase activity of MlaFEDB was significantly increased to an average of 0.024 μM ATP s^−1^ at 500 μM ATP; whilst this represents a significant increase in ATP hydrolysis, it remains a significantly lower rate than is exhibited by MlaFEDB in detergent micelles however (Figure 8). Together these results suggest that PL environment is essential in regulating MlaFEDB ATPase activity, and that when constrained in a bilayer MlaFEDB requires the interaction of MlaC-apo to drive ATP hydrolysis, further highlighting the importance of the MlaC PL bound state in defining PL movement.

**Figure 8.**
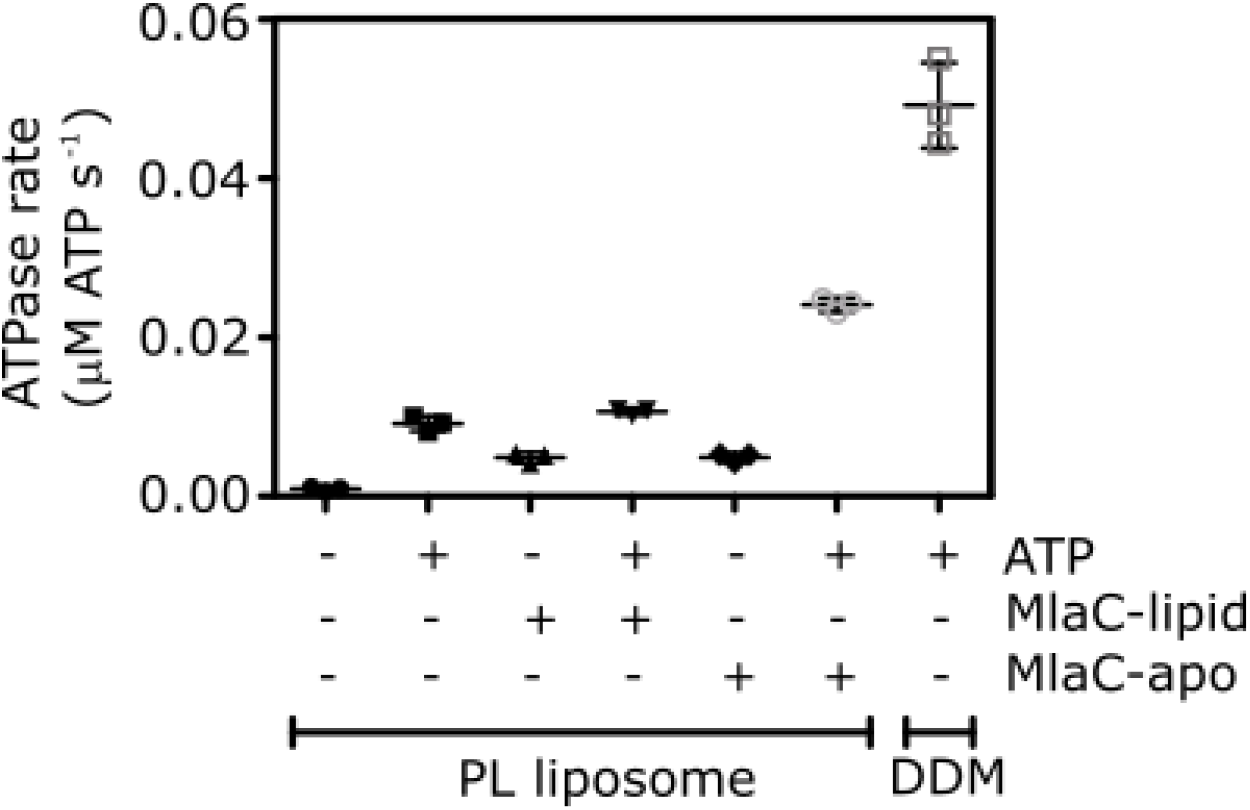
MlaFEDB ATPase activity upregulated in the presence of MlaC-apo. Enzyme coupled ATPase assay of MlaFEDB *E. coli* polar lipid proteoliposomes in the presence and absence of 500μM ATP, MlaC-apo and MlaC-lipid. Shown for comparison is the equivalent ATPase rate for DDM solubilised MlaFEDB.

### ATP hydrolysis has no obvious effect on phospholipid transfer

QCM-D, FTIR and TLC allow for a more in depth analysis of the role of ATP hydrolysis in PL transport. Using QCM-D and measuring mass changes during interaction with MlaC (apo or lipid) in the presence or absence of ATP we were able to probe changes in rate. Strangely no obvious differences in the rate of PL loss from the surface were noted (Figure 6A & B). The mass changes observed on addition of either MlaC-apo or MlaC-lipid remained comparable irrespective of the presence or absence of ATP suggesting that the removal of PL from the surface was ATP independent. We have already shown that ATPase activity is still present following binding to Cu^2+^ NTA beads and reconstitution in to a PL bilayer suggesting that surface tethering should not impact function (Figure S9). However to confirm this in the absence of 6xHis tethering, we used FTIR (Figure 7C). Similar results were noted, the rate of PL movement from MlaFEDB to MlaC-apo occurred at the same rate irrespective of the presence of ATP and again no PL movement was noted from MlaC-lipid to MlaFEDB either in the presence or absence of ATP. Furthermore, using our TLC transport assay, neither the absence of ATP nor the presence of the non-hydrolysable ATP analogue, ATP-γS, impacted on the observed transport of PLs from MlaFEDB to MlaC-apo. For both conditions (Figure 5D & E), similar PL transport was noted between MlaFEDB and MlaC-apo whilst no transport was noted in the retrograde direction. These results combined with the observation that ATPase activity is upregulated in the presence of MlaC-apo suggest that the ATPase activity of MlaFEDB is not directly related to the PL export process but has an alternate function within the Mla pathway.

## Discussion

It is generally thought that the Mla pathway maintains OM lipid asymmetry by removing PLs from the outer leaflet of the OM then shuttling them across the periplasmic space via MlaC to the inner membrane (Malinverni and Silhavy 2009). Here, MlaFEDB, an ABC transporter, through utilising ATP, was hypothesised to insert those PLs in to the IM (Malinverni and Silhavy 2009, Thong, Ercan et al. 2016). The ability of MlaC to act as a PL shuttle between the IM and OM is reliant upon its reversible transition between PL bound and PL free states, here for the first time we demonstrate the ability of MlaC to form this PL free condition. Furthermore, through crystallographic studies we have determined its structure to a resolution of 2.25 Å, and show that during the transition between PL bound and PL free states, MlaC experiences global structural changes, involving a unique β-sheet pivot mechanism. Furthermore, by probing MlaC PL binding states, we have shown that, rather than MlaFEDB functioning to take PLs from MlaC and insert them in to the inner membrane, it appears to do the opposite and extracts PLs from the IM and passes them to MlaC and does so in an ATP independent manner.

Our elucidation of the structure of MlaC-apo has shed light on its ability to accommodate various PL groups. Its binding mechanism, involving a unique β-sheet pivot, which results in a closing of the PL binding cavity in the MlaC-apo condition is supportive of molecular dynamics investigations performed by Huang *et al*. (2016) who suggested that the size of the PL binding pocket of MlaC from both *R. solanacearum* and *Acinetobacter baumannii* would be dramatically reduced upon PL release (Huang, Miao et al. 2016). Simulations performed herein using experimentally determined MlaC structures from three different species have confirmed these previous results and suggest that this mechanism of closure is likely to be widespread across Gram-negative bacteria. Moreover, as there appears to be no obvious restriction on the movement of the β-sheet, it suggests the ability to pivot further, opening the cavity more to occupy PLs such as cardiolipin as suggested by Ekiert *et al*. (2017) (5uwb) possibly explaining MlaC’s ability to accommodate multiple lipid types (Malinverni and Silhavy 2009). To this end, molecular modelling simulations have suggested that MlaC may be able to fine-tune the topography of its binding site to accommodate different PLs (Huang, Miao et al. 2016). This is an important concept to consider when thinking about MlaC as a PL shuttle, as it would perhaps allow a tailoring of which PLs were transported depending on necessity at the OM.

Understanding the molecular mechanisms of PL transport from the IM to the OM represents one of the major challenges surrounding bacterial PL homeostasis (Henderson, Zimmerman et al. 2016). Whilst it has been well established that a host of PLs can move in a retrograde fashion from the OM to the IM (Jones and Osborn 1977), it remains unclear as to how PLs travel in the opposite direction. It has been suggested that retrograde PL transport is in part driven by the action of the OM MlaA-OmpF/C system (Abellon-Ruiz, Kaptan et al. 2017). In fact, Thong *et al*. (2016) proposed that the MlaFEDB driven hydrolysis of ATP may provide the necessary energy to indirectly remove PLs from the OM against their concentration gradient. Here we provide evidence to the contrary, and suggest that MlaFEDB can directly transport PLs from the IM and transfer them to MlaC. Furthermore, we observed that the apo form of MlaC is solely required for this unique movement of PLs, no utilisation of ATP is required. These findings represent a novel mechanism for MlaFEDB driven anterograde PL transport, and allows us to think of MlaC as a potential PL-shuttle, funneling PLs toward the OM MlaA complex. This is feasible considering how dominant mutations in MlaA result in both increased levels of PLs at the OM and a concurrent deterioration of the IM (Sutterlin, Shi et al. 2016). In addition, Malinverni & Silhavy (2009) were unable to follow the Mla dependent movement of PLs out of the OM, again suggestive of a system inclined toward anterograde PL transport.

Through a combination of QCMD, FTIR and TLC we have demonstrated that the movement of PLs from MlaFEDB to MlaC occurs independently of ATP hydrolysis, with transport occurring at comparable rates regardless of MlaFEDB ATPase inhibition or the presence of 500 μM ATP. This is feasible considering the transport of PLs from the IM to the OM would involve movement down a concentration gradient. Furthermore, Sutterlin *et al*. (2016) noted that even in response to ATP synthase inhibition, dominant MlaA mutations continue to promote PL flow away from the IM, again suggesting passivity in this mechanism.

Whilst it has been suggested that ATP hydrolysis may facilitate PL transfer from MlaC to MlaFEDB (Ekiert, Bhabha et al. 2017), our evidence suggests otherwise and as such the role of the MlaFEDB ATPase remains ambiguous. Therefore, elucidating how ATPase activity is coupled to PL transport represents an important area for future study. Considering the function of MlaFEDB at the IM, it is possible that its ATPase activity may be linked to potential flippase ability, with MlaC binding promoting MlaFEDB dependent ATP hydrolysis and a subsequent flipping of PLs between inner and outer leaflets, which may be significant in maintaining the integrity of the IM. In fact, a similar ABC transporter, MsbA, has been shown to function in a comparable manner at the IM, where it acts to flip LPS and PLs from the inner to the outer leaflet in an ATP dependent fashion (Doerrler and Raetz 2002, Eckford and Sharom 2010). In addition, we have shown MlaFEDB dependent ATP hydrolysis to be highly regulated in terms of PL environment and putative MlaC interactions, with ATPase activity dramatically increasing exclusively in the presence of MlaC-apo. This is unsurprising considering the ATPase activity of other well-characterized ABC transporters is intrinsically linked to the binding of their corresponding substrate-binding proteins (Liu, Liu et al. 1997, Chen, Sharma et al. 2001). Moreover, this would avoid redundant ATP hydrolysis, allowing for potential coupling of MlaFEDB flippase ATPase activity to the process of MlaC PL loading.

The model we present here, in which MlaFEDB is involved in PL transport out of the IM (Figure 9), opens up the notion of MlaC acting as a soluble, periplasmic PL store. Whilst we have focused on MlaFEDB driven PL transport, it is possible that in situations in which both MlaFEDB and MlaA are present this process would not occur in such a unidirectional fashion. As such, *in vivo* MlaC may function as a PL sensor, with its overall lipid state defining the directionality of transport. This mechanism would require exclusively affinity driven interactions between MlaC and the different Mla complexes at both the IM and OM, which may be possible considering the passive nature of PL transfer from MlaFEDB to MlaC-apo. Therefore, if MlaC is involved in bidirectional PL transport, the inability of MlaC-lipid to deliver PLs into a bilayer via MlaFEDB would signal towards an as yet unknown mechanism for MlaC PL recycling, which given the number of MCE protein complexes involved in as of yet poorly understood PL transport processes, represents a viable concept (Ekiert, Bhabha et al. 2017).

**Figure 9.**
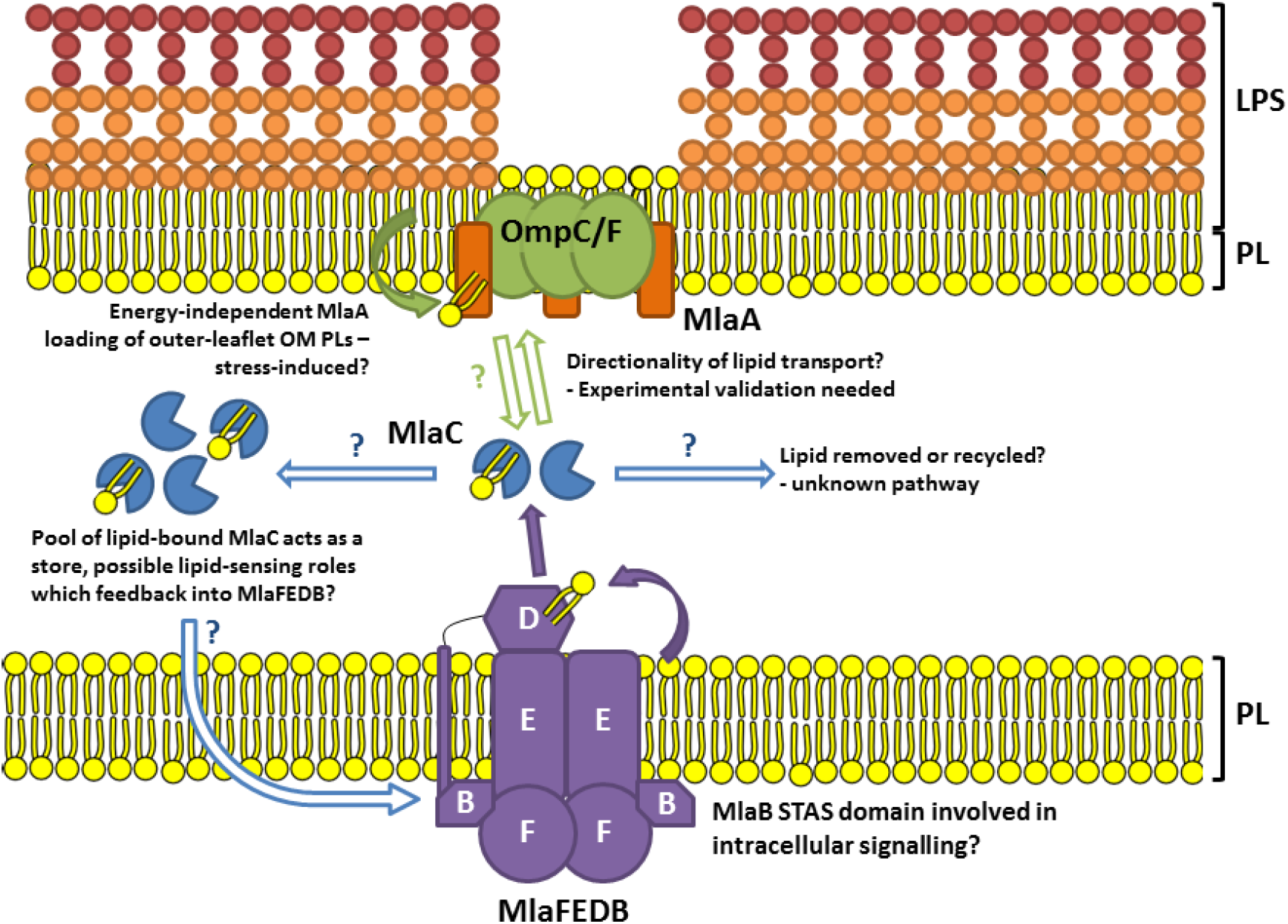
Schematic summarising questions for the mechanisms of the Mla system arising from this study. Experimental evidence strongly suggests that MlaFEDB is capable of, and shows a preference for, transferring PL to MlaC. Further work is needed to fully elucidate the functions of each of the Mla components, which likely reach beyond that of the classically theorised retrograde PL trafficking.

## Conclusions

In summary, this study provides compelling evidence for PL export from the inner membrane to the periplasmic space and suggests rather than retrograde transport, the Mla pathway may be involved in anterograde transport to the bacterial outer membrane and that the export process appears to occur in an ATP independent manner. What role the ATPase activity of MlaFEDB plays in this process still remains to be seen but we speculate that it may function as a flippase within the inner membrane, maintaining the symmetry of the membrane and preventing the curvature that would be expected if the export process occurred only from a single leaflet.

## Acknowledgements

We thank D. Ekiert and G. Bhabha for discussion and kindly providing Mla plasmid constructs. We thank HWB-NMR at the University of Birmingham for providing open access to their Wellcome Trust-funded NMR equipment. This research was supported by BBSRC grant BB/P009840/1 (TJK & GWH) and Wellcome Trust grant 208400/Z/17/Z. TJP acknowledges use of the Iridis high-performance computing resources at the University of Southampton.

## Supplementary Figure Legends

**Supplementary Figure 1.**
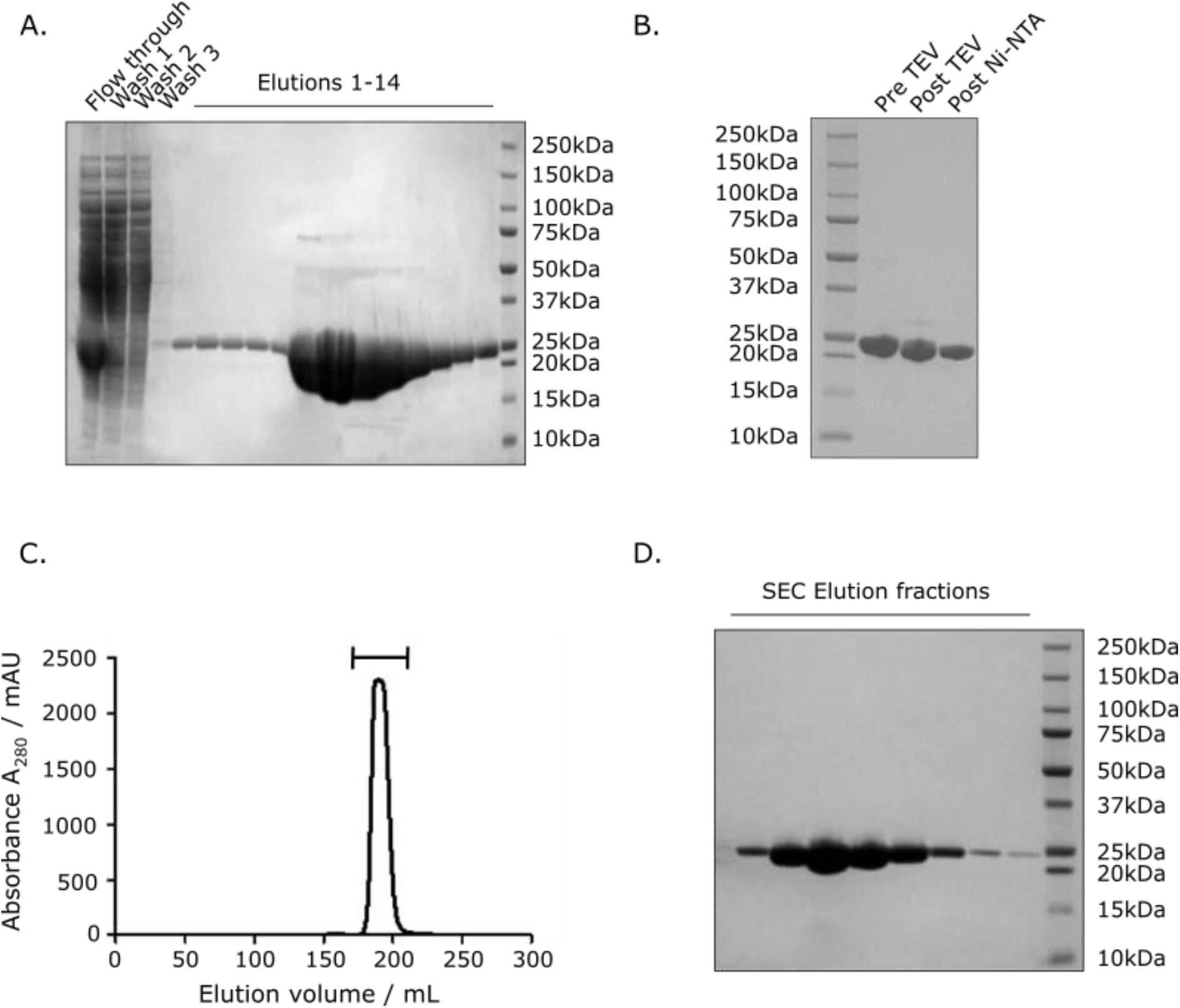
Purification of MlaC. A) SDS-PAGE showing the Ni-NTA purification of MlaC. B) SDS-PAGE of MlaC pre/post TEV cleavage and subsequent cleanup following passage through Ni-NTA column. C) S75 Size exclusion chromatogram showing the presence of a single uniform peak consistent with monomeric MlaC. D) SDS-PAGE showing post size exclusion fractions containing purified MlaC.

**Supplementary Figure 2.**
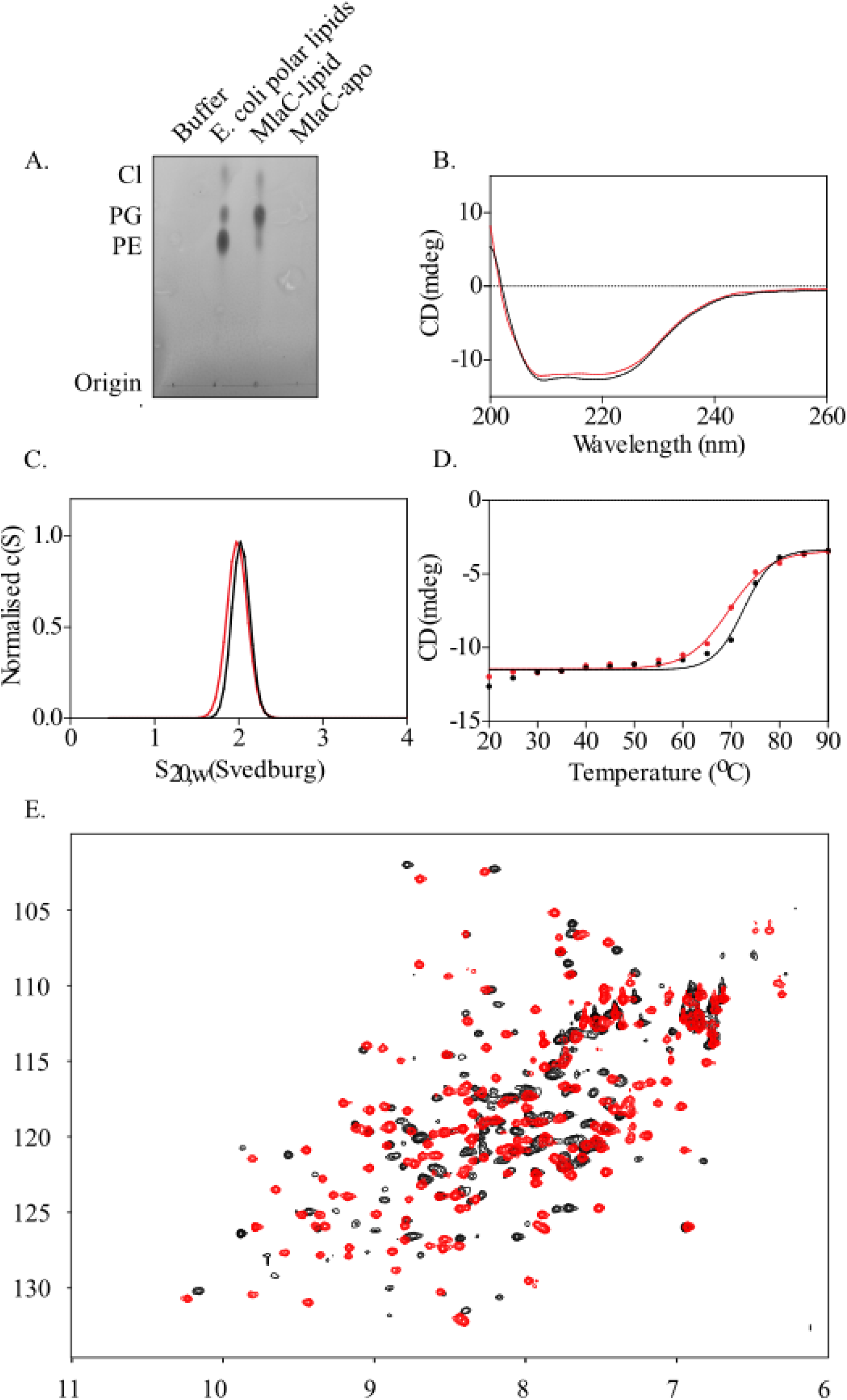
Biophysical characterisation of MlaC-apo. Biophysical comparison of PL bound and PL free MlaC. A) Thin layer chromatogram confirming detergent washing of MlaC-lipid leads to removal of PL and formation of MlaC-apo. B) Circular dichroism spectra of MlaC-lipid (black) and MlaC-apo (red) confirming MlaC-apo maintains fold on PL removal. C) Analytical ultracentrifugation sedimentation velocity analysis of MlaC-lipid (black) and MlaC-apo (red) confirming no significant change in mass on PL loss. D) Circular dichroism thermal melt recorded at 208 nm of MlaC-lipid (black) and MlaC-apo (red) showing a 3°C reduction in thermal stability on PL loss. E) ^1^H^15^N-HSQC spectra of 0.3mM ^15^N-labelled MlaC in the presence (black) and absence (red) of PL highlighting significant changes in chemical shift perturbations throughout MlaC on PL removal.

**Supplementary Figure 3.**
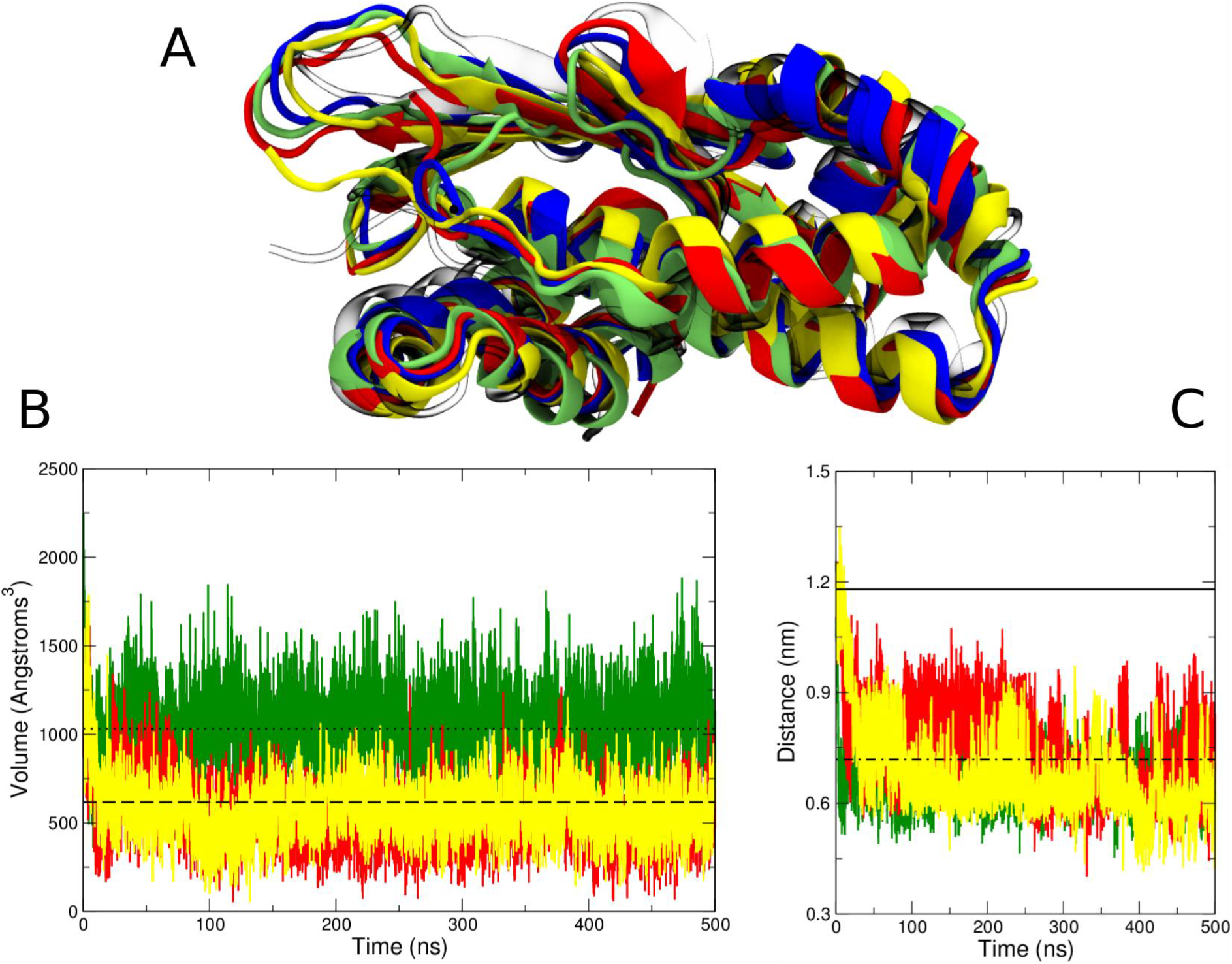
Demonstration of *E. coli* MlaC closure upon phospholipid removal. A) Superimposed structures of the MlaC from the end (500 ns) of three representative simulations performed using structure 5uwa with the PL removed, one from each of the force fields used (AMBER99SBMR-ILDN red, CHARMM36 yellow and GROMOS 54A7 green). The simulation starting structure is also shown (transparent white), as is the MlaC-apo structure (blue). B) Changes in cavity volume during the same simulations as shown in A, with the same colour scheme applied. The cavity volume from the MlaC-apo structure is also shown (dashed all-atom protein representation, dotted united-atom protein representation). C) Minimum distances between Cα atoms of residues 99-109 (located within helix α5) and residues 164-171 (final turn of the β-sheet) during the same three representative simulations. The distances in the starting structure (solid line) and MlaC-apo structure (dotted and dashed line) are also shown for reference. The shorter distance in the MlaC-apo structure is indicative of the closure at the mouth of the cavity.

**Supplementary Figure 4.**
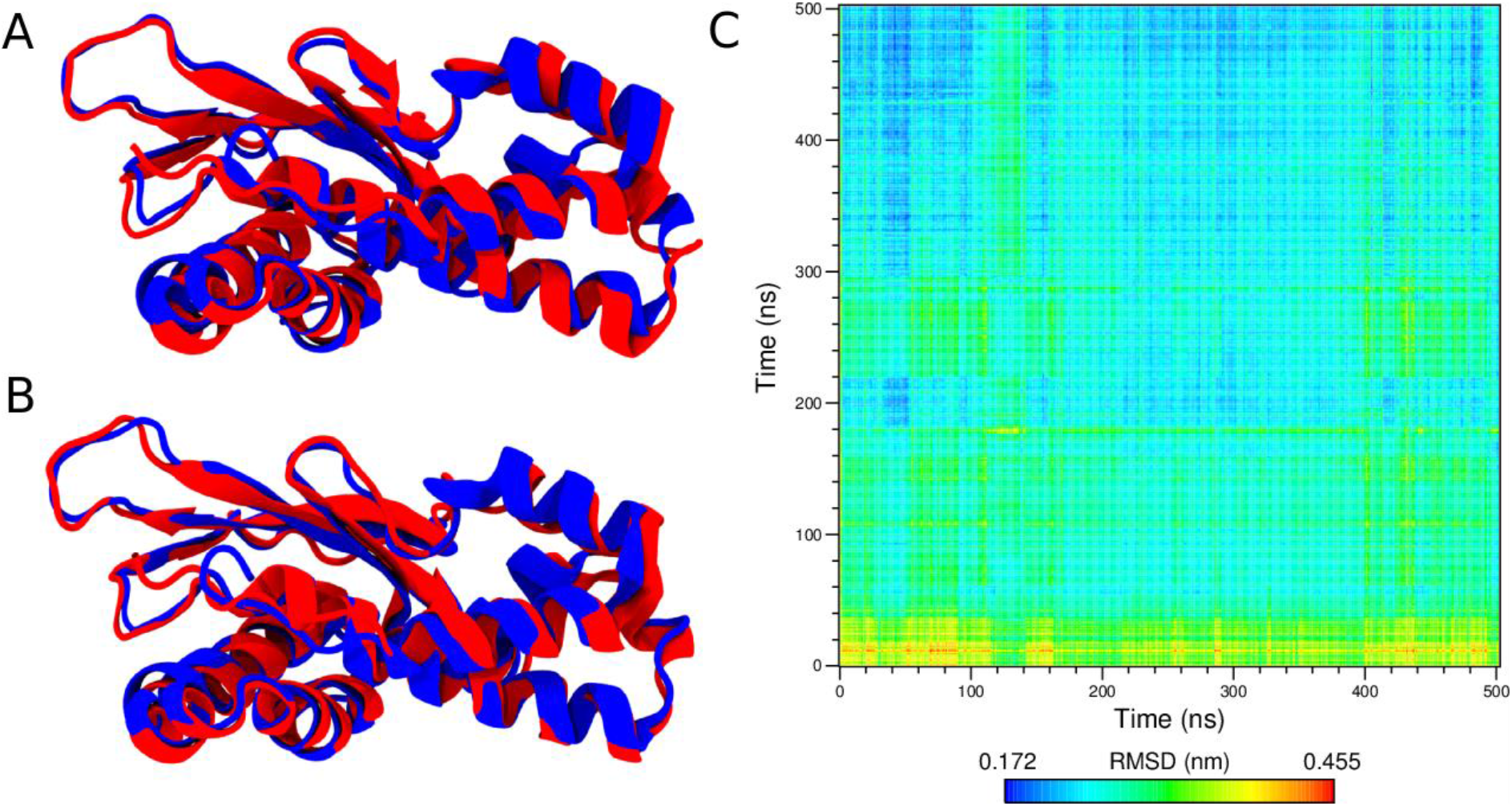
MlaC-apo conformational stability and similarity to the simulated apo structures. A) and B) Superimposed structures at the start (blue) and end (red) of the standard molecular dynamics simulations performed using the MlaC-apo structure. C) Comparison of the MlaC protein structure from one of these MlaC-apo simulations (x-axis) to the protein structure from one of the apo simulations performed using the 5uwa starting structure with the CHARMM36 force field (y-axis). A smaller root mean squared deviation (RMSD) indicates closer similarity. After the initial collapse of the cavity in the simulation initiated from the open structure (5uwa), the structures adopt similar apo conformations despite the use of different force fields.

**Supplementary Figure 5.**
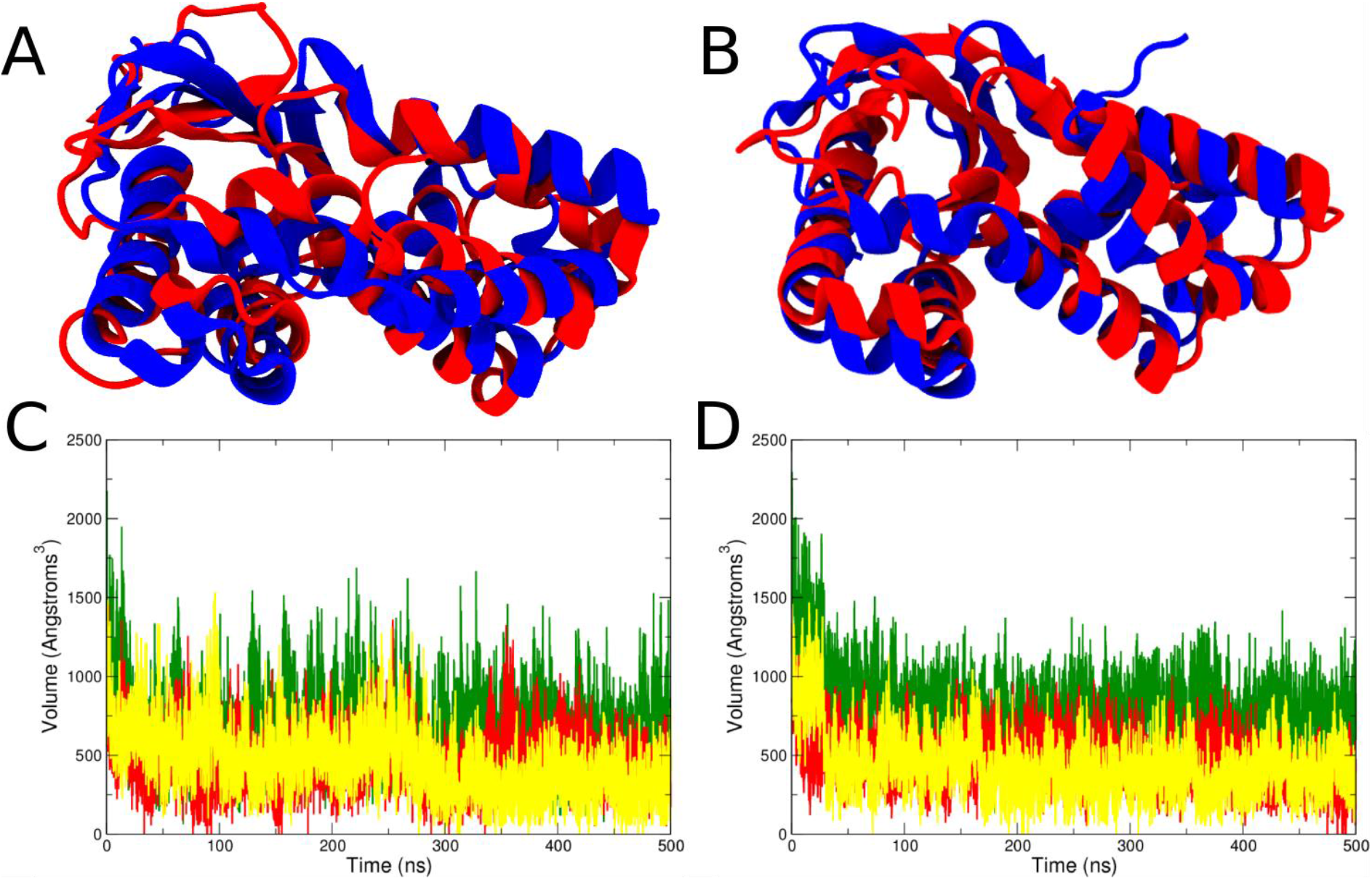
*R. solanacearum* and *P. putida* MlaC closure upon phospholipid removal. A) and B) Superimposed structures of the starting (blue) and final (red) conformations from representative A) *R. solanacearum* and B) *P. putida* apo simulations. We note that 8 unstructured residues at the C-terminus of the *P. putida* MlaC structure have been removed from the images for clarity. C) and D) Changes in cavity volume during three representative simulations using different force fields of MlaC-apo from *R solanacearum* (C) and *P. putida* (D). The colour scheme in C) and D) follows that of Figure S4.

**Supplementary Figure 6.**
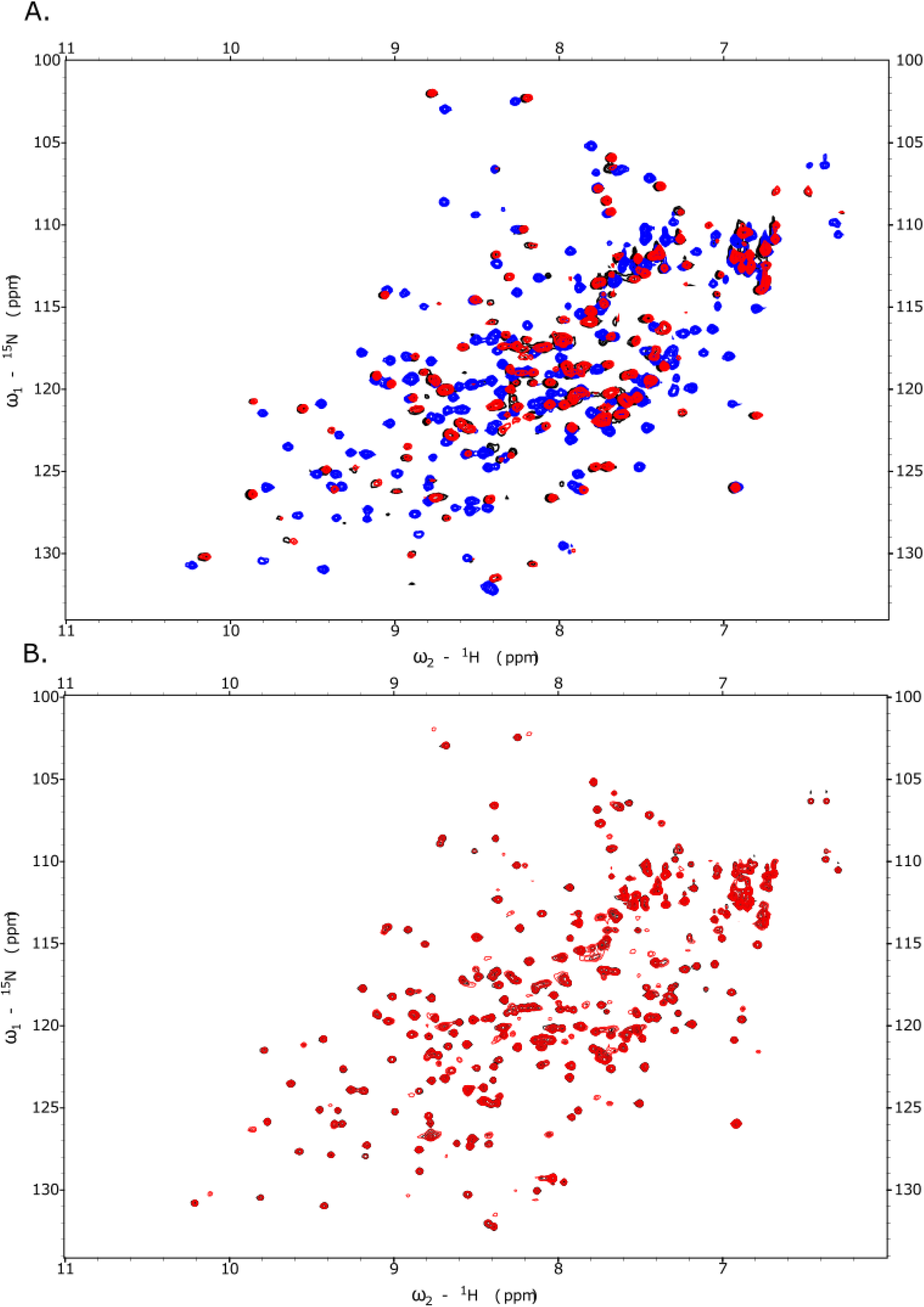
NMR analysis of phospholipid binding. A) ^1^H^15^N HSQC spectra of 0.3mM ^15^N-MlaC in its PL bound form (Black), after detergent washing to remove PL and produce MlaC-apo (Blue), and MlaC-apo following 30 min incubation with PL bound MlaD and subsequent SEC purification (red). B) ^1^H^15^N HSQC spectra of 0.2mM MlaC-apo (Black) and MlaC-apo following incubation with 2mg·mL-1 *E. coli* polar lipids for 18 hours (red).

**Supplementary Figure 7.**
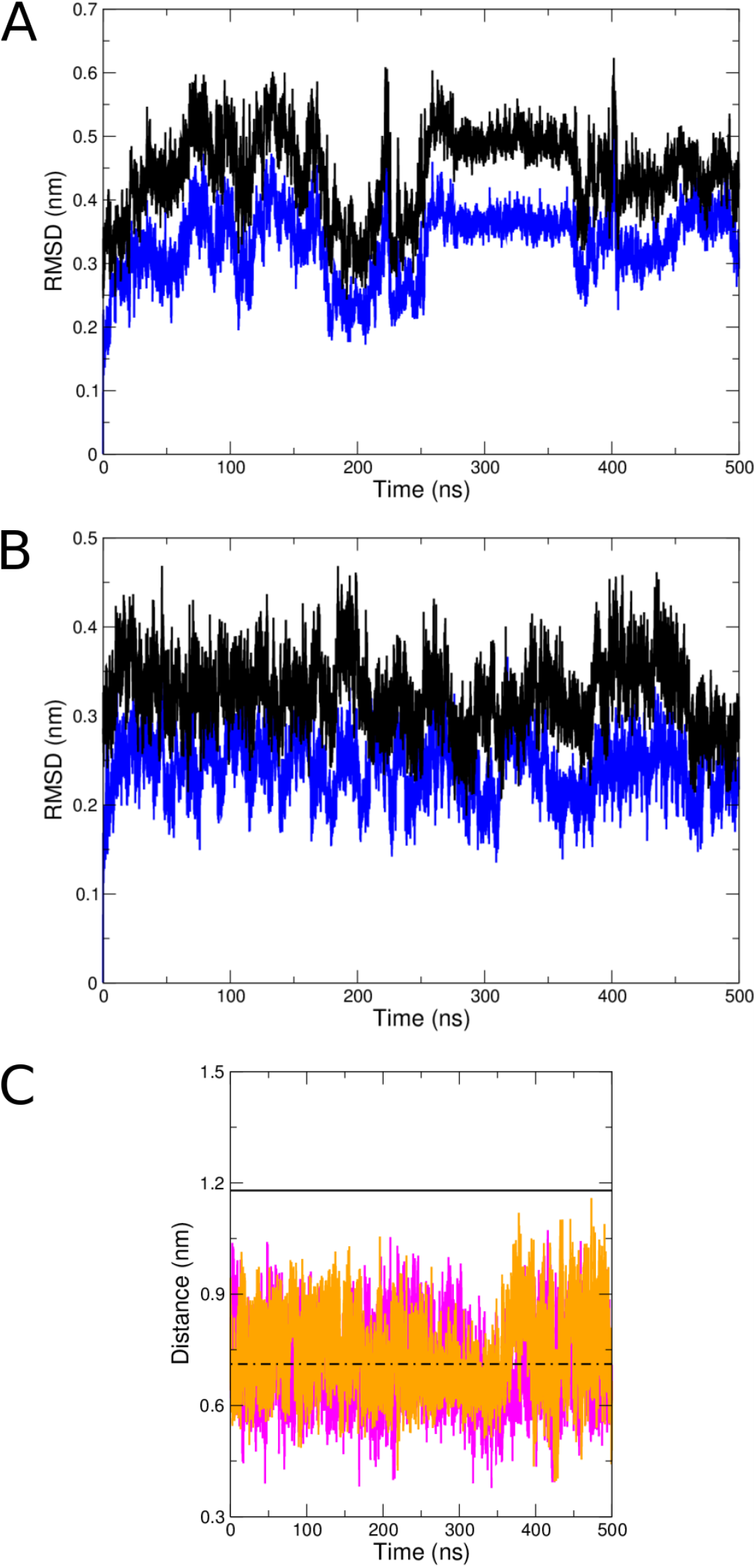
Enhanced sampling simulations fail to sample the MlaC-lipid structure. Total protein RMSD during one high temperature (A) and one scaled-down interaction (B) simulation of the MlaC-apo structure. In these figures the RMSD to the MlaC-apo structure is shown in blue and the RMSD to the PL bound 5uwa structure is shown in black. The overall similarity of the protein in PL bound and apo states can be seen in the relative similarity of the RMSDs. C) Minimum distances between Cα atoms of residues 99-109 (located within helix α5) and residues 164-171 (final turn of the β-sheet) during one high temperature (orange) and one scaled interaction (pink) simulation. As per Figure S4, the distances in the 5uwa and MlaC-apo structure are also shown for reference. In both high temperature and scaled interaction simulations the protein conformations remain closer to that of MlaC-apo with the cavity closed.

**Supplementary Figure 8.**
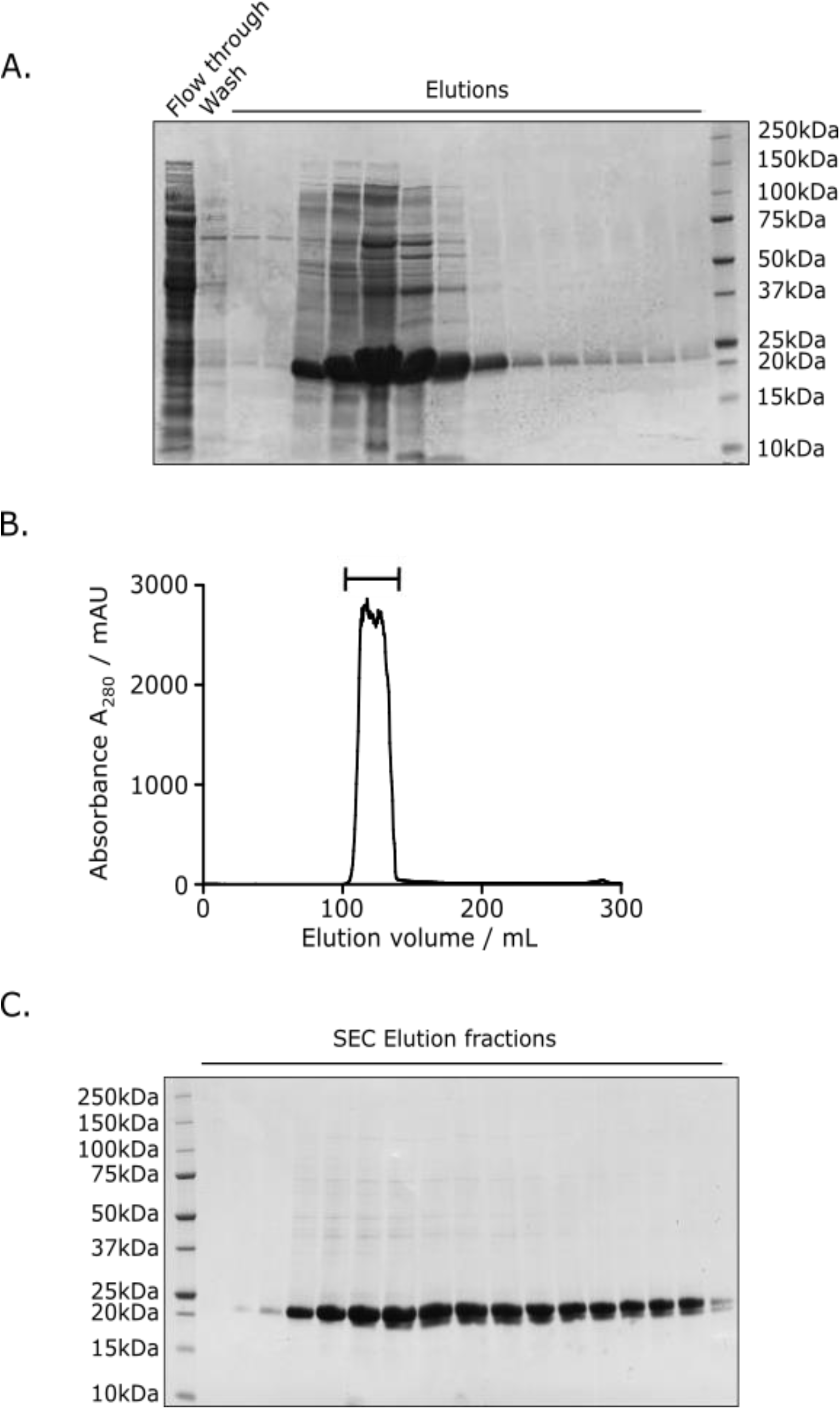
Purification of MlaD. SDS-PAGE showing the Ni-NTA purification of MlaD. B) S75 Size exclusion chromatogram showing the presence of a single uniform peak consistent with hexameric MlaD. D) SDS-PAGE showing post size exclusion fractions containing purified MlaD.

**Supplementary Figure 9.**
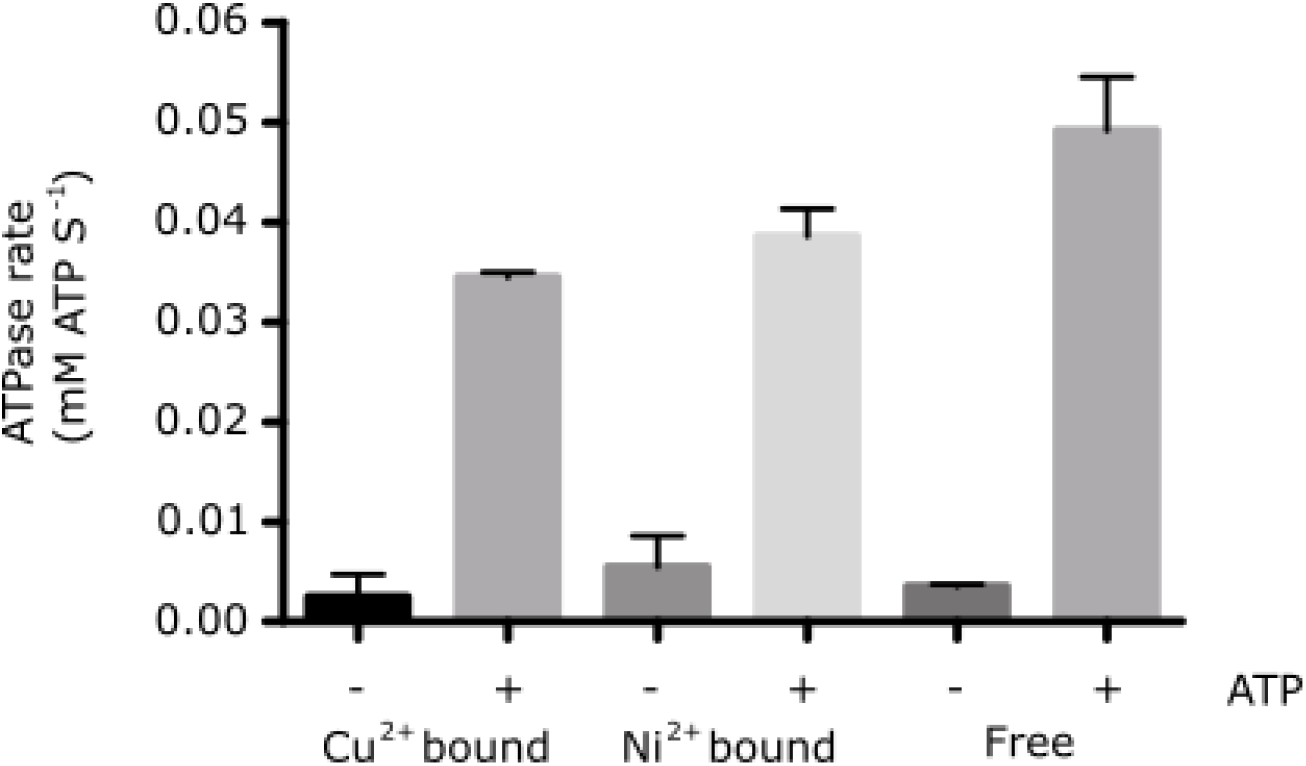
Resin tethered MlaFEDB shows ATPase activity. Enzyme coupled ATPase assay of detergent solubilised MlaFEDB in the presence and absence of 500μM ATP either free in solution or bound to either Ni^2+^ or Cu^2+^-NTA beads.

**Supplementary Figure 10.**
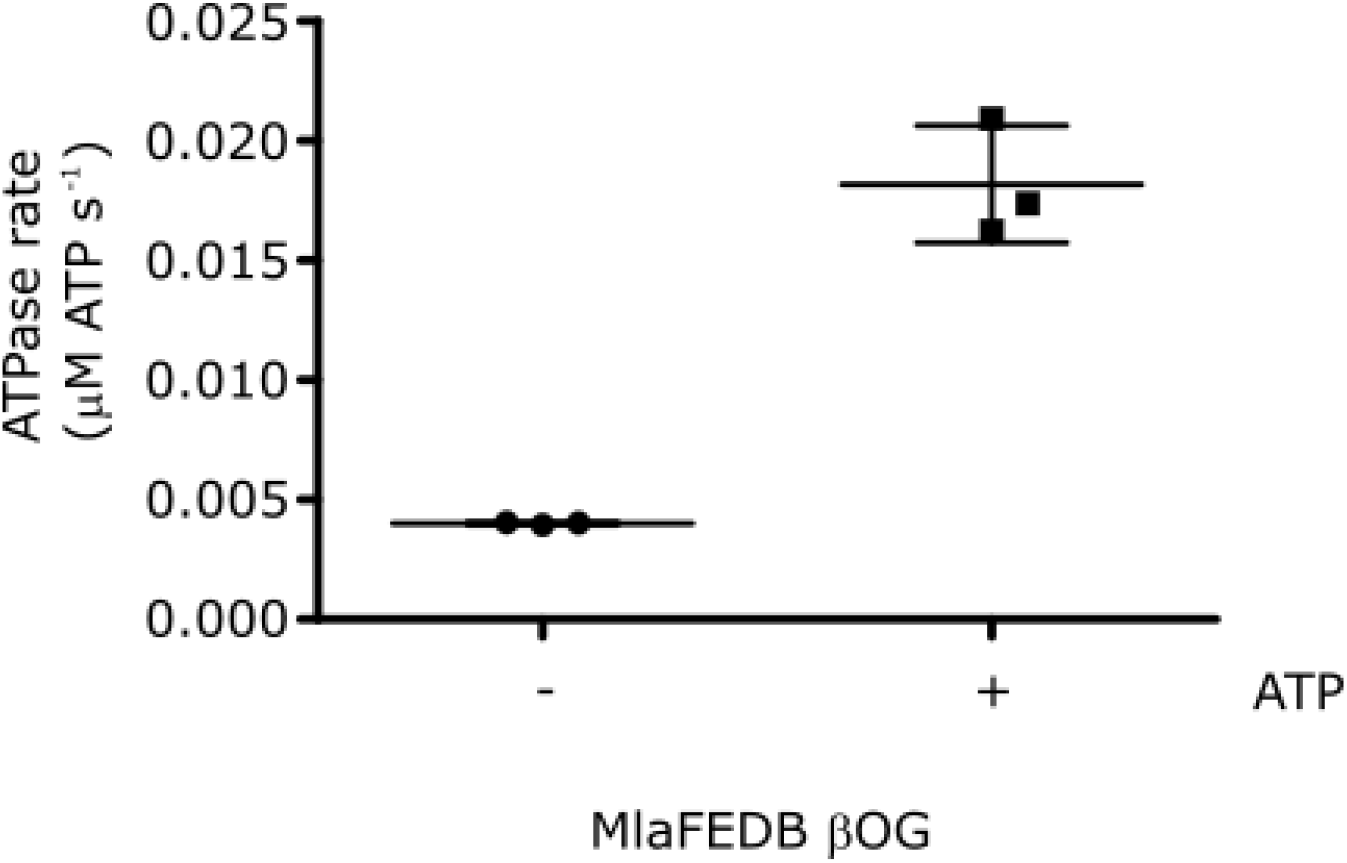
MlaFEDB shows activity in β-octyl glucoside. Enzyme coupled ATPase assay of MlaFEDB (0.1μM) after detergent exchange in to 25mM β-octyl glucoside in the presence and absence of 500μM ATP.

**Supplementary Figure 11.**
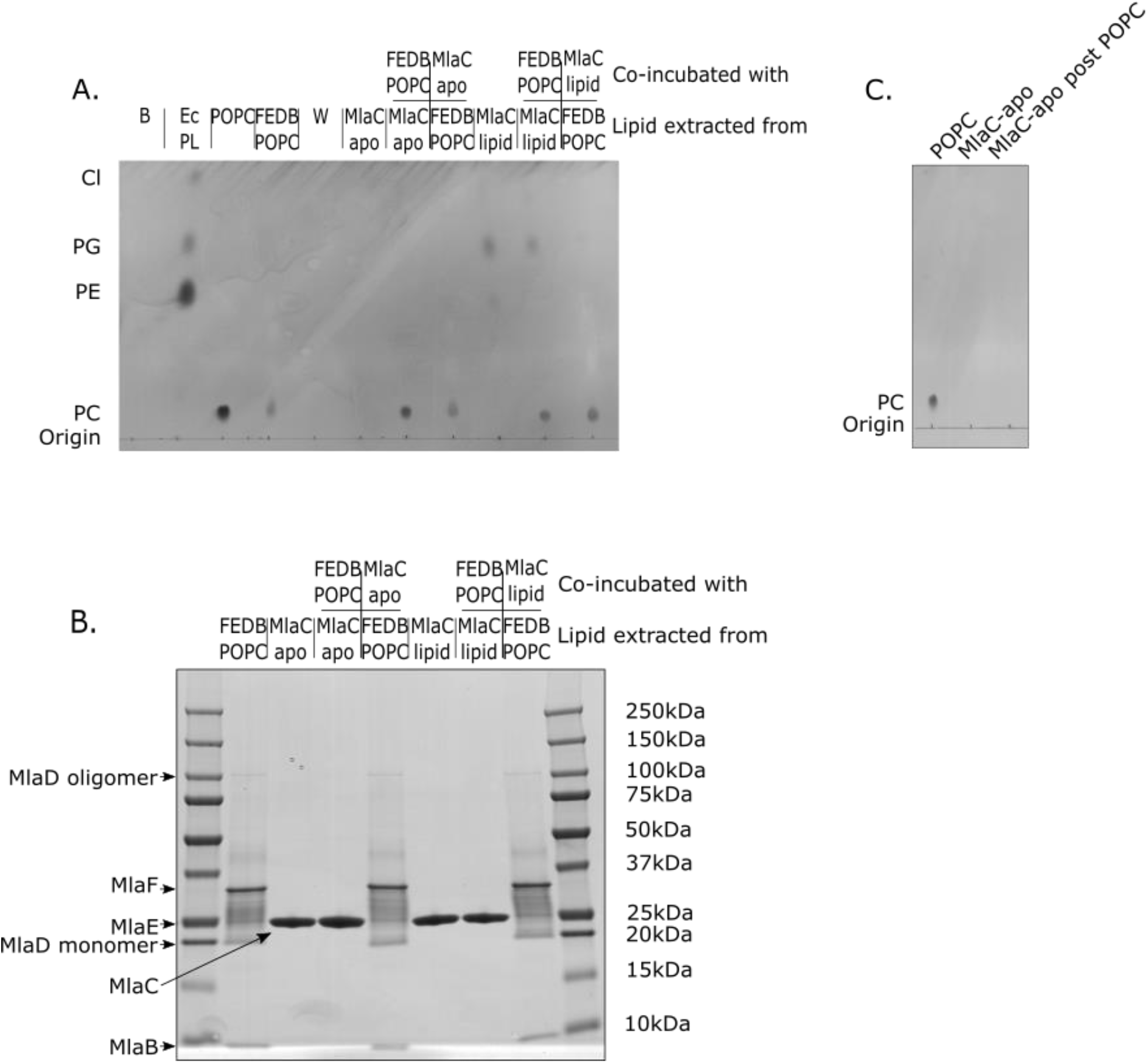
POPC has no detrimental effects on MlaFEDB function. TLC showing the movement of PL between MlaFEDB and MlaC following reconstitution of MlaFEDB in to POPC liposomes. B – Buffer; Ec PL – *E. coli* Polar lipids; POPC - 1-palmitoyl-2-oleoyl-sn-glycero-3-phosphocholine; MlaFEDB POPC – MlaFEDB reconstituted in POPC liposomes; W – Wash following binding of MlaFEDB-POPC to a Ni-NTA column. B) SDS-PAGE of the samples shown in A) confirming separation of the various species following incubation. C) TLC of MlaC-apo and MlaC-apo following incubation with POPC (MlaC-apo post POPC) showing no addition of PL to MlaC occurs.

## Supplementary Table Legends

**Supplementary Table 1.**
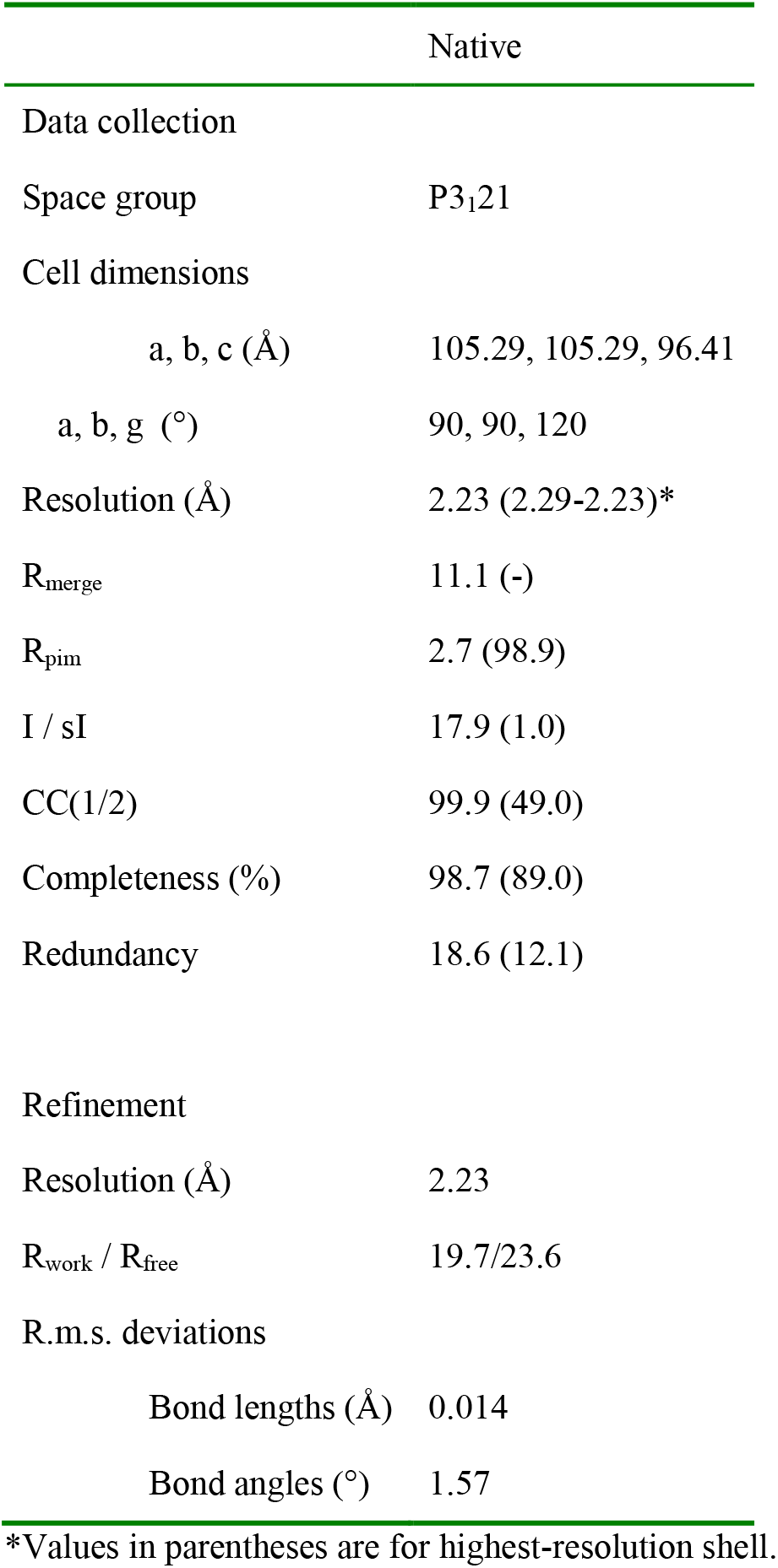
X-ray diffraction data collection and refinement statistics

**Supplementary Table 2.** *E. coli* MlaC pocket closure upon phospholipid removal quantified by solvent accessible surface area (SASA). SASA at the start and averaged over the final 100 ns of all the apo simulations initiated from the 5uwa structure with the PL removed. The standard deviation of the average over the final 100 ns is also provided in parentheses. The consistent reduction in the SASA is indicative of the closure of the cavity in all of these simulations.

| **Simulation ID** | **SASA Start (nm<sup>2</sup>)** | **Average SASA Final 100 ns (nm<sup>2</sup>)** |
| --- | --- | --- |
| AMBER 1 | 114.81 | 108.01 (1.92) |
| AMBER 2 | 114.98 | 107.07 (2.24) |
| AMBER 3 | 114.99 | 105.08 (1.91) |
| AMBER 4 | 115.45 | 104.84 (2.07) |
| AMBER 5 | 115.11 | 104.64 (1.84) |
| CHARMM 1 | 114.26 | 107.60 (2.12) |
| CHARMM 2 | 114.29 | 104.82 (2.04) |
| CHARMM 3 | 114.10 | 103.48 (1.84) |
| CHARMM 4 | 114.23 | 105.81 (1.89) |
| CHARMM 5 | 113.73 | 105.05 (1.55) |
| GROMOS 1 | 119.19 | 106.81 (2.20) |
| GROMOS 2 | 117.48 | 107.96 (2.70) |
| GROMOS 3 | 118.18 | 106.40 (2.18) |
| GROMOS 4 | 120.19 | 105.22 (2.27) |
| GROMOS 5 | 119.08 | 107.88 (2.06) |

